# Homeostatic Synaptic Plasticity Rescues Neural Coding Reliability

**DOI:** 10.1101/2021.12.06.471391

**Authors:** Eyal Rozenfeld, Nadine Ehmann, Julia E. Manoim, Robert J. Kittel, Moshe Parnas

## Abstract

To survive, animals must recognize reoccurring stimuli. A key requirement for repeated identification of stimuli is reliable representation by the neural code on each encounter. Synaptic transmission underlies neural codes propagation between brain regions. A hallmark of chemical synapses is their plasticity, which enables signal transfer to be modified in an activity-dependent manner. Despite many decades of intense research on synapses, it remains unclear how the plastic features of synaptic transmission can maintain reliable neural coding. By studying the olfactory system of *Drosophila melanogaster*, we aimed to obtain a deeper mechanistic understanding of how synaptic function shapes neural coding reliability in the live, behaving animal. We show that the properties of the active zone (AZ), the presynaptic site of neurotransmitter release, are critical for generating a reliable neural code. Reducing neurotransmitter release probability specifically at AZs of olfactory sensory neurons disrupted both neural coding and behavioral reliability. Strikingly, these defects were rescued within a day by target-specific synaptic plasticity, whereby a homeostatic increase in the number of AZs compensated the drop in release probability. These findings demonstrate an important role for synaptic plasticity in maintaining neural coding reliability and are of pathophysiological interest by uncovering an elegant mechanism through which the neural circuitry can counterbalance perturbations.

## Introduction

Animals encounter the same stimuli repeatedly and are able to recognize them over and over again. An essential requirement for the repeated identification of a stimulus is that its representation by the neural code will be reliably reproduced on each occasion. Two major approaches are used to describe neural coding^1^. The first is rate coding, where the firing rate of action potentials over a period of time is employed as the neural code. The second is temporal coding, where the precise timing and the precise pattern of action potentials is utilized for coding. Often, a combination of the two strategies is used as the coin of information for neural coding: the firing rate within a relatively small time window along with changes in firing rate over time.

Coding of an external stimulus by the nervous system begins at the sensory neurons. The code then undergoes transformations as it passes across synaptic contacts and propagates to higher brain regions. Chemical synaptic transmission is the major mode of fast information transfer between neurons. The response of a postsynaptic neuron depends on neurotransmitter release from synaptic vesicles (SVs) fusing with the plasma membrane at the presynaptic active zone (AZ). SV fusion, in turn, is regulated by the complex interplay of various specialized AZ proteins, which determine neurotransmitter release sites and control release probability^2–4^. The molecular mechanisms of SV fusion are highly dynamic and can be modified on timescales ranging from milliseconds to days. Such presynaptic plasticity involves e.g. changes in Ca^2+^ signals, SV availability, and activity-dependent modulations of the release machinery^5–9^. Importantly, synapses are also under homeostatic control. Homeostatic synaptic plasticity describes a phenomenon whereby pre- or postsynaptic adjustments maintain synaptic stability^10–13^. This evolutionarily conserved process rebalances destabilizing perturbations and preserves functionality in a changing environment. Correspondingly, dysfunctional homeostatic synaptic plasticity has been implicated in several neurological diseases^14,15^. Considering the many facets of synaptic plasticity, the question arises how the reliability of the neural code is affected by dynamic changes in neurotransmission^16–20^.

The *Drosophila* olfactory system is highly amenable to genetic manipulations and is suitable to examine how synaptic function and plasticity affect neural code reliability. In flies, odors activate olfactory receptor neurons (ORNs). In general, each ORN expresses a single odorant receptor gene^21–23^ (but note ref.^24^). Approximately 20-40 ORNs expressing the same receptor send their axons to a single glomerulus in the antennal lobe (AL)^25–27^. Approximately 2-5 second-order projection neurons (PNs), in turn, send their dendrites to a single glomerulus^28^ and each PN samples all arriving ORNs^29^. The synaptic connection between an individual ORN and PN harbors ~10-30 release sites (scaling with glomerulus size) possessing a high average release probability (~0.75)^30^. The AL also contains multi-glomeruli inhibitory GABAergic and glutamatergic local neurons (iLNs) and excitatory cholinergic LNs (eLNs)^31–34^. The ORN-PN synapse is mainly modulated by the GABAergic iLNs^34,35^. Thus, the *Drosophila* ORN-PN connection is a system with a reliable synapse, which can be genetically modified and examined in the context of physiological odor stimuli.

By performing *in vivo* whole-cell patch-clamp recordings from over 1300 neurons, we show that knocking down the AZ Ca^2+^ channel Cacophony (Cac) specifically in ORNs decreases the reliability of the neural code in the postsynaptic PNs to physiological odor stimuli. Our results demonstrate that interfering with SV release probability from ORN AZs affects the PN neural code in three ways. First, the initial transient phase of the odor response, when PNs reach their highest firing rates, becomes more variable due to reduced recruitment of iLNs. Second, the onset of the odor response is delayed and more variable. Third, the temporal dynamics of the odor response become less reliable; this third result does not involve any circuit motifs and results from monosynaptic effects. In line with its ethological relevance, reducing the reliability of neural coding leads to a reduction in the behavioral ability to correctly classify an olfactory stimulus. Surprisingly, however, we find that decreased expression of Cac at ORN AZs affects neural code reliability only at high stimulus intensities. This differential effect is due to homeostatic synaptic plasticity, which sets in within a day. Thereby, the formation of additional AZs compensates the reduced neurotransmitter release probability and rescues the coding reliability of PNs. Thus, our work uncovers a new role for homeostatic synaptic plasticity in maintaining stable neural coding and reliable behavior.

## Results

### ORN SVs release probability affects olfactory coding reliability at high odor intensities

In order to investigate how molecular properties of the AZ influence the olfactory neural code, we recorded postsynaptic PN responses to odor presentation with *in vivo* whole-cell current-clamp recordings (Figure 1A). We used RNA interference (RNAi) in ORNs to knock down Cac, the pore-forming subunit of the *Drosophila* Cav2-type Ca^2+^ channel, which mediates Ca^2+^-influx at the AZ^36,37^. Since different odors elicit different firing rates and temporal dynamics in ORNs^38^ we tested PN responses to five different odors (Isopenthyl acetate, linalool, 2-heptanone, ethyl acetate and isobutyl acetate) and used high odor concentrations to minimize variations in ORN responses^39^. To examine stimulus-evoked response reliability we recorded PN responses to 10 applications of each odor (Figure 1A-C and Figure S1). *Orco-GAL4* was used to drive *UAS-Cac^RNAi^* specifically in ORNs and *GH146-QF*, which labels approximately 60% of PNs^40^, was used to drive *QUAS-GFP* for targeted patch-clamp recordings. We confirmed RNAi efficiency for Cac by performing *in vivo* Ca^2+^ imaging via two-photon microscopy during odor stimulation. *Orco-GAL4* was used to drive *UAS-Cac^RNAi^* along with *UAS-GCaMP6f*. As expected, Ca^2+^ signals were significantly reduced in ORN terminals when Cac was targeted by RNAi (Figure S2A). In addition, direct staining against Cac demonstrated reduced Cac levels in *Cac^RNAi^* flies (Figure S2B). *Cac^RNAi^* significantly reduced odor-induced PN activity (Figure 1B, C and Figure S2C), illustrating that SV release from ORNs was decreased, though not completely abolished.

**Figure 1:**
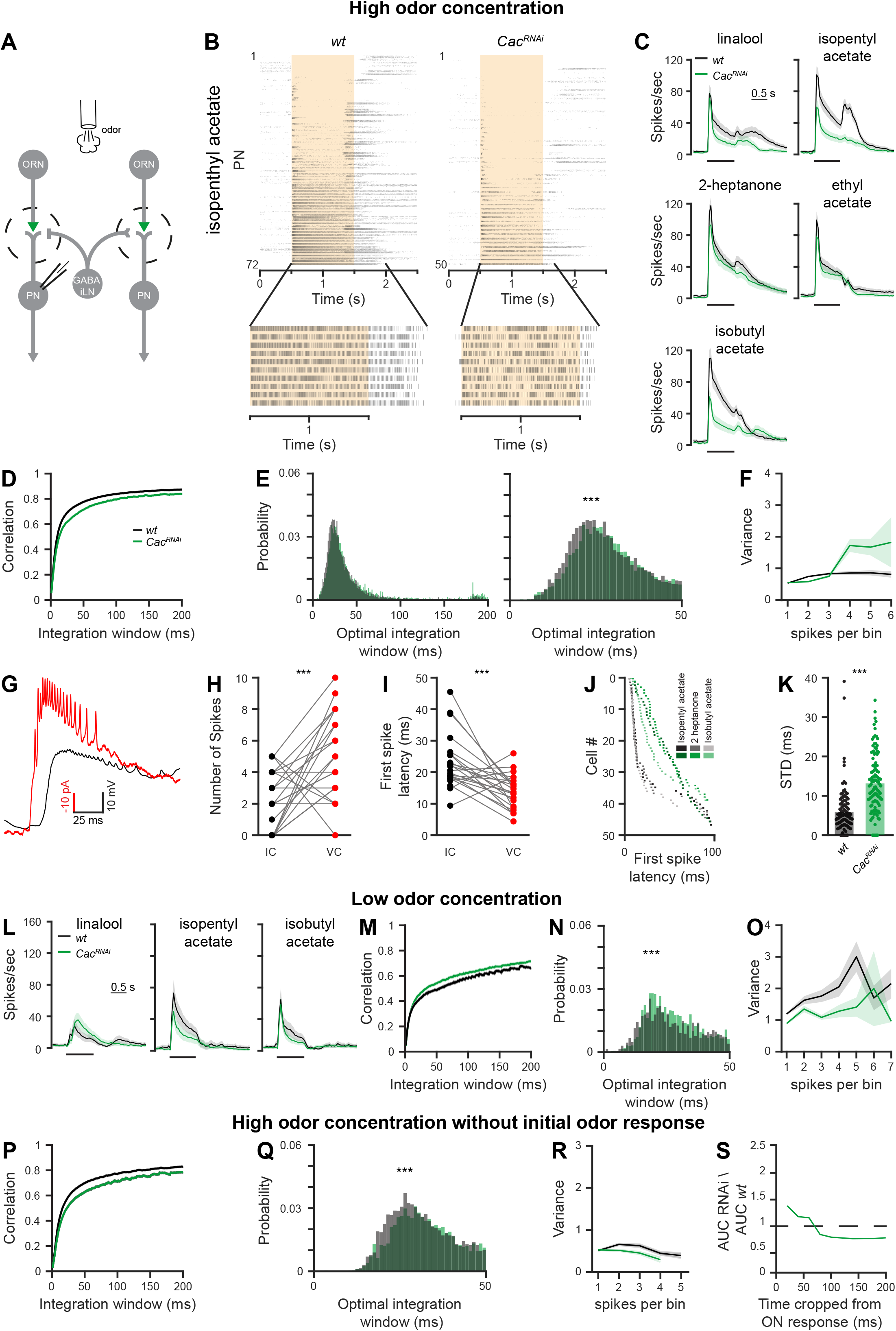
Reducing SV release probability reduces coding reliability only at high stimulus intensities. **A.** Experimental scheme. *UAS-Cac^RNAi^* was expressed in ORNs using *Orco-GAL4* and whole-cell patch clamp recordings were made from PNs labeled by the *GH146-QF* driver line. PN odor response were measured for high odor concentration in 2-4 days old flies. **B.** *Top*, Raster plot of PN population odor responses to isopenthyl acetate (a final odor dilution of 5X10^−2^ was used) in *wt* flies (left, n=72 flies) and *Cac^RNAi^* in ORNs (right, n=50 flies). Each neuron was presented with 10 repetitions of the olfactory stimulus (1 s). *Bottom*, the 10 repetitions of a single PN are presented. The shaded area indicates the odor stimulus. **C.** Peristimulus time histogram (PSTH) of PN population responses to five odors examined as indicated (shaded areas represent SEM, odor pulse is labeled with a black bar) obtained from recordings as in panel A. Spike trains were binned using a 50 ms time bin. Knockdown Cac (green) in ORNs resulted in decreased PN odor responses. A final odor dilution of 5X10^−2^ was used (n=48-72 flies). *Orco-GAL4* was used to drive the RNAi constructs and *GH146-QF* drove *QUAS-GFP*. **D.** Temporal reliability analysis. Pairwise correlations for each odor-neuron combination were pooled across all odors for data in Figure S1. Non-overlapping windows from 1 to 200 ms were used. *Cac^RNAi^* in ORNs reduces correlation values. **E.** *Left*, the curve saturation point (see methods) was calculated for each odor-neuron combination and pooled across all odors for data in Figure S1. *Right*, Left curves are presented at a larger scale. *Cac^RNAi^* in ORNs shifts the optimal temporal integration window of PNs as evident by a larger integration window spread (left) and a peak shift of ~10 ms (right). **F.** Firing-rate reliability analysis for data in Figure S1. Spike trains were binned using 20 ms windows. Spike count and variability were calculated for each bin and pooled across neurons, odors, and time. Increased rate variability is observed for high firing rates. **G.** Example traces of recordings performed in current clamp (IC, black) and in voltage clamp (VC, red). Note, the VC trace is inverted for presentation purposes. **H.** PN firing-rate during the first 25 ms of the response to isopentyl acetate in current clamp (IC, black) vs. voltage clamp (VC, red). Each dot represents an individual fly (n=22). The VC configuration enabled detection of more action potentials. **I.** First spike latency of PNs in response to isopentyl acetate in IC (black) vs. VC (red). Dots represent individual flies (n=22). **J.** First spike latency of PNs in response to the indicated odors for *wt* and *Cac^RNAi^* flies (n=50). Each dot represent the mean first spike latency for 10 trials of a given neuron. Data were obtained in VC configuration and PNs that did not spike within 100ms after stimulus onset were omitted. **K.** First spike jitter of PN odor responses, pooled across all odors, for *wt* and *Cac^RNAi^* flies (n=50). **L.** PSTH of PN population responses to three low concentration odors examined as indicated (shaded areas represent SEM, odor pulse is labeled with a black bar). Spike trains were binned using a 50 ms time bin. Knockdown Cac (green) in ORNs did not affect PN odor responses. A final odor dilution of 5X10^−4^ was used (n=50 flies). *Orco-GAL4* was used to drive the RNAi constructs and *GH146-QF* drove *QUAS-GFP*. **M, N.** Temporal reliability analysis (as in panels D, E) for data in panel L. *Cac^RNAi^* in ORNs did not reduce correlation values. **O.** Firing-rate reliability analysis (as in panel F) for data in panel L. *Cac^RNAi^* in ORNs did not increase firing rate variability. **P, Q.** Temporal reliability analysis (as in panels D, E) for data in Figure S1. Removing the first 200 ms of the odor response leads to an overall decrease in correlation. A shift of the peak of the curve is observed for the sustained odor response in *Cac^RNAi^* flies. **R.** Firing-rate reliability analysis (as in panel F) for data in Figure S1 without the first 200 ms of the odor response. Contrary to the increased variation observed for the entire odor response (Figure 2C), *Cac^RNAi^* in ORNs did not increase the variability for the sustained odor response although high firing-rates were still obtained. **S.** The ratio between the area under the curve (AUC) of the rate code variability in *Cac^RNAi^* relative to *wt*. The initial response was progressively cropped from the analysis. The difference in rate code variation of *wt* and *Cac^RNAi^* flies gradually decreases over the first 200 ms of the odor response. For all panels *** p<0.001, see table S1 for statistical analysis.

A number of different methods can be used to examine neural code reliability. Correlation-based measurement of coding reliability examines the reliability of the temporal dynamics of neural activity by analyzing the activity time-series as a single entity^41^. Thus, this analysis examines the pattern of the response over time, but does not consider the magnitude of the response. Therefore, correlation measurements do not account for variations in firing rate at a specific time point (Figure S3A, C). Another measure of temporal code reliability is the latency of the first spike following odor presentation, which was show to encode odor identify and intensity^42^. The standard deviation of response magnitude examines response reliability for a given time point or a given firing rate and does not account for the temporal dynamics (Figure S3B, C)^43^.

At high odor concentration, correlation analysis using spike trains that were integrated at increasingly wider temporal windows (Figure S3A), revealed that for wild type *(wt)* flies the response reliability (correlation value) became saturated at ~20 ms (Figure 1D, E), consistent with previous reports^44,45^. Knockdown of Cac decreased overall correlation values and increased the temporal integration window required to reach saturation (Figure 1D, E and Figure S4A, B). This demonstrates that molecular perturbations of ORN AZs decrease the temporal reliability of PN responses. Standard deviations of PN firing rate, as a function of the mean firing rate during a 20 ms time window, also revealed a decrease in PN coding reliability in *Cac^RNAi^* flies (Figure 1F and Figure S4C) with PNs showing increased variability especially at high firing rates (Figure 1F).

PN somata, which are accessed during patch-clamp recording, are distant from the action potential initiation site^46,47^. As a result, action potentials recorded at the soma in current-clamp mode are very small, often reaching only 5 mV in amplitude^46^. Such small action potentials are frequently not identified during the rising phase of the odor response. To better detect the first action potential, indicating the response onset, we repeated this set of experiments in voltage-clamp mode. Indeed, voltage-clamp recordings enabled detection of more action potentials in the first 25 ms of the odor response and the latency of the first action potential was shorter (Figure 1G-I). As with the other neural coding reliability measurements, this analysis revealed that knocking down Cac resulted in lower reliability, reflected by both increased latency and increased standard deviation of the first action potential (Figure 1J, K and Figure S5).

High odor concentrations elicit high firing rates in ORNs that may reach up to 300 Hz^48^. To examine if the effect of *Cac^RNAi^* persists when ORNs fire at lower rates, we repeated the above experiments using a weaker odor intensity. Interestingly, we found that under these conditions, response magnitude and coding reliability by PNs was not decreased by reduced Cac expression (Figure 1L-O). Taken together, our data show that reducing SV release from ORNs affects the reliability of the postsynaptic odor response. However, this effect is only evident during the presentation of a strong olfactory stimulus.

The above results suggest that high-frequency synaptic transmission is most sensitive to manipulations of ORN AZs, consistent with previous reports^49^. High firing rates usually occur at the beginning of the odor response within the first 200 ms (Figure 1B, C). Thus, we examined the effects of *Cac^RNAi^* during the presentation of a strong olfactory stimulus without the initial phase of the odor response. The initial phase of the odor response is characterized by a strong increase in firing rate that is highly correlated. Thus, as expected, removing the first 200 ms of the odor response from the analysis resulted in an overall decrease in correlation (Figure 1P compare to Figure 1D), which was further reduced by *Cac^RNAi^* (Figure 1P, Q). In contrast, the increased standard deviation of PN firing rates that was observed upon *Cac^RNAi^* was now completely abolished. No difference in the standard deviation was observed between *wt* and RNAi flies even though high firing rates were still obtained in the sustained phase of the odor response (Figure 1R). The effect of *Cac^RNAi^* was not equally distributed throughout the initial phase of the odor response; rather it was most notable when the PNs reached their peak firing rate, approximately 20ms after the response onset (Figure 1S). These results suggest that two different mechanisms underlie the reduced response reliability of PNs, one affecting the reliability of the structure of the temporal firing rate dynamics and the other affecting the reliability of the absolute firing rate magnitude.

### Knockdown of Cac reduces release probability and increases synaptic latency and jitter

Reducing Cac expression in ORNs significantly reduced PN spiking (Figure 1). Nevertheless, PNs maintained a substantial response to odors, demonstrating that synaptic transmission from ORNs to PNs remained principally functional. Cac is a key regulator of SV release probability at *Drosophila* neuromuscular AZs. Its functional disruption in a temperature-sensitive mutant, RNAi-mediated knockdown, and impaired AZ channel clustering all lead to greatly decreased eEPSC (evoked excitatory postsynaptic current) amplitudes at the NMJ, especially at low stimulation frequencies^36,37,50^. We were therefore intrigued by the observation that *Cac^RNAi^* affected PN coding reliability only at high and not at low ORN firing rates. To better understand this phenomenon, we characterized functional properties of the ORN-PN synapse in *wt* and *Cac^RNAi^* animals (Figure 2A). Surprisingly, the average eEPSC amplitude of *Cac^RNAi^* did not differ from *wt* (Figure 2B). However, Cac knockdown led to increased paired-pulse facilitation at short inter-pulse intervals (Figure 2C, D), consistent with a decrease in SV release probability upon reduced expression of AZ Ca^2+^ channels^50^. In line with a drop in release probability, *Cac^RNAi^* synapses also displayed significantly elevated synaptic delay (eEPSC latency; Figure 2E), which was accompanied by an increase in the jitter of eEPSC (Figure 2F). Interestingly, both parameters progressively increased with stimulation frequency in the transgenic animals (Figure 2E, F). Within our sampling range (1, 10, 20, 60 Hz), the effect of presynaptic *Cac^RNAi^* on these temporal parameters was most pronounced at 60 Hz.

**Figure 2:**
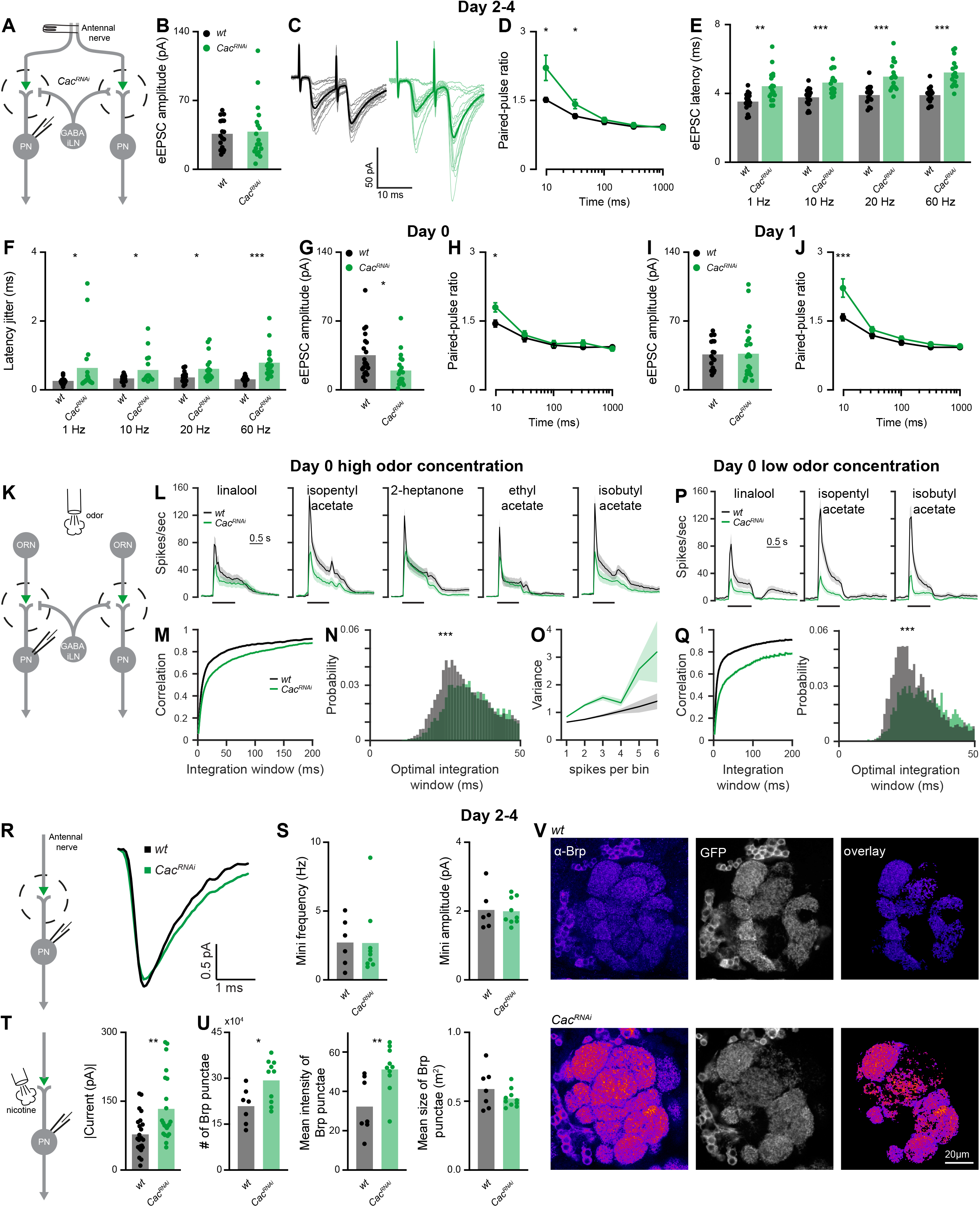
Presynaptic Cac knockdown induces homeostatic plasticity. **A.** Experimental scheme. *UAS-Cac^RNAi^* was expressed in ORNs using *Orco-GAL4* and whole-cell patch clamp recordings were made from PNs labeled by the *GH146-QF* driver line. The antennal nerve was stimulated with a suction electrode. **B.** The average eEPSC amplitude was unaltered by *Cac^RNAi^* for 2-4 days old flies (1 Hz stimulation frequency; *Orco-Gal4* driver, *wt* n=17 and *Cac^RNAi^* n=18 flies). **C.** Examples traces of eEPSC evoked by paired pulse stimulation with an inter-pulse interval of 10ms. **D.** Quantification of the paired-pulse ratio at different inter-stimulus intervals for 2-4 days old flies (10ms, 30ms, 100ms, 300ms and 1000ms). *Cac^RNAi^* significantly increased paired-pulse facilitation at short inter-pulse intervals (*wt*, n=20-22; *Cac^RNAi^*, n=24-28 flies). Error bars represent SEM. **E, F.** Average eEPSC latencies (**E**) and average eEPSC jitter (**F**) at 1, 10, 20, and 60 Hz stimulation for *wt* (n=17) and *Cac^RNAi^* (n=18) flies. A significant increase was observed for all tested stimulation frequencies (except of jitter at 1 Hz). **G.** The average eEPSC amplitude was reduced by *Cac^RNAi^* for 0 day old flies (1 Hz stimulation frequency; *Orco-Gal4* driver, *wt* n=20 and *Cac^RNAi^* n=19 flies). **H.** Quantification of the paired-pulse ratio at different inter-stimulus intervals for 0 day old flies (10ms, 30ms, 100ms, 300ms and 1000ms). *Cac^RNAi^* significantly increased paired-pulse facilitation at short inter-pulse intervals (*wt*, n=20-21; *Cac^RNAi^*, n=19 flies). Error bars represent SEM. **I.** The average eEPSC amplitude was unaltered by *Cac^RNAi^* for 1 day old flies (1 Hz stimulation frequency; *Orco-Gal4* driver, *wt* n=17 and *Cac^RNAi^* n=18 flies). **J.** Quantification of the paired-pulse ratio at different inter-stimulus intervals for 1 day old flies (10ms, 30ms, 100ms, 300ms and 1000ms). *Cac^RNAi^* significantly increased paired-pulse facilitation at short inter-pulse intervals (*wt*, n=20-22; *Cac^RNAi^*, n=22 flies). Error bars represent SEM. **K.** Experimental scheme. *UAS-Cac^RNAi^* was expressed in ORNs using *Orco-GAL4* and whole-cell patch clamp recordings were made from PNs labeled by the *GH146-QF* driver line. PN odor response were measured for both high and low odor concentration in 0 days old flies. **L.** PSTH of PN population responses to five high concentration odors examined as indicated (shaded areas represent SEM, odor pulse is labeled with a black bar) for 0 day old flies. Spike trains were binned using a 50 ms time bin. Knockdown Cac (green) in ORNs resulted in decreased PN odor responses. A final odor dilution of 5X10^−2^ was used (n=50-57 flies). *Orco-GAL4* was used to drive the RNAi constructs and *GH146-QF* drove *QUAS-GFP*. **M, N.** Temporal reliability analysis (as in Figure 1 D, E) for data in panel L. *Cac^RNAi^* in ORNs reduces correlation values. **O.** Firing-rate reliability analysis (as in Figure 1 F) for data in panel L. *Cac^RNAi^* in ORNs increased firing rate variability. **P.** PSTH of PN population responses to three low concentration odors examined as indicated (shaded areas represent SEM, odor pulse is labeled with a black bar) for 0 day old flies. Spike trains were binned using a 50 ms time bin. Knockdown Cac (green) in ORNs resulted in decreased PN odor responses. A final odor dilution of 5X10^−4^ was used (n=49-50 flies). *Orco-GAL4* was used to drive the RNAi constructs and *GH146-QF* drove *QUAS-GFP*. **Q.** Temporal reliability analysis (as in Figure 1 D, E) for data in panel P. *Cac^RNAi^* in ORNs reduces correlation values. **R.** *Left*, Experimental scheme. Spontaneously occurring single-vesicle fusions (minis) were recorded at PNs in 2-4 days old flies. *Right*, Representative traces of averaged spontaneous minis in PNs of *wt* (black) and *Cac^RNAi^* in ORNs (green). **S.** Analysis of mini frequency (left) and amplitude (right) does not show significant differences between genotypes. (*wt*, n=6; *Cac^RNAi^*, n=9 flies). **T.** *Left*, Experimental scheme. Nicotine (100μM) was injected to the AL and currents were measured in PNs. *Right*, Analysis of the current magnitude in response to a nicotininc puff. *Cac^RNAi^* in ORNs increases PN response to the cholinergic agonist nicotine, indicating an increase in postsynaptic receptor fields. *Orco-GAL4* was used to drive the RNAi construct and *GH146-QF* drove *QUAS-GFP (wt*, n=23; *Cac^RNAi^*, n=22 flies). **U.** Analysis of Brp in ORN pre-synapses shows a significant increase in number and fluorescence intensity of Brp punctae following Cac knock-down, while the size of Brp points is unaffected. (*wt*, n=7; *Cac^RNAi^*, n=10 flies). **V.** Example confocal images of an individual plane through the antennal lobe in control (upper panel; *orco- Gal4/+; GH-146-QF,QUAS-GFP*/+) and Cac^RNAi^ flies (lower panel*; orco-Gal4/+; GH146-QF,QUAS-GFP*/*UAS-Cac^RNAi^*) stained against Brp (fire). To restrict the analysis of Brp to excitatory PN post-synapses, the Brp signal was overlayed with masks generated from imaging endogenous GFP driven via GH146 (GFP, grey; overlay). Scale bar: 20 μm. For all panels, * p<0.05, ** p<0.01, *** p<0.001, see table S1 for statistical analysis.

Taken together, these results suggest a homeostatic mechanism at the ORN-PN synapse, which compensates for the drop in release probability caused by presynaptic Cac knockdown to maintain normal eEPSC amplitudes. However, this compensation does not cover the temporal properties of synaptic transmission. The reduced release probability of *Cac^RNAi^* ORNs generates longer synaptic latencies with larger eEPSC jitter, consistent with increased first spike latencies and jitter during odor application. Thus, while the ORN-PN synapse is surprisingly resilient to Ca^2+^ channel perturbations, Cac knockdown delays synaptic transmission and reduces the temporal precision of synaptic signaling.

### A homeostatic increase in AZ number and synaptic strength compensates for the drop in release probability

So far, all experiments were carried out on 2-4 day old flies. To obtain more information on the homeostatic regulation of the ORN-PN connection, we next addressed the time course of the synaptic adjustment. At day 0 (less than 24 h post pupal eclosion), *Cac^RNAi^* caused a reduction in both release probability and eEPSC amplitudes (Figure 2G, H). Moreover, *Cac^RNAi^* synapses responded in only 19 out of 40 cases irrespective of stimulus intensity. In contrast, at day 1 (24-48 h post pupal eclosion), eEPSC amplitudes were already elevated to *wt* levels (Figure 2I, J) and the transmission success rate reached 100%. When presented with a high odor concentration at day 0, *Cac^RNAi^* flies displayed decreased firing rates and reduced coding reliability compared to *wt* (Figure 2K-O) and to 2-4 days old flies (Figures 1C-F and Table S1). However, when a low odor concentration was applied, both firing rates and neural code reliability remained severely impaired (Figure 2P, Q), contrary to the compensation observed in 2-4 days old flies (Figure 1L-O). Thus, homeostatic synaptic plasticity took place within the first day post eclosion to rescue neural activity and coding reliability at low stimulus intensity but failed to do so at high odor concentrations.

Next, we turned to the mechanism underlying the homeostatic synaptic change responsible for maintaining normal eEPSC amplitudes. In principle, the drop in release probability caused by *Cac^RNAi^* could be counterbalanced by an increase in the number of presynaptic release sites or an increase in quantal size. Quantal size describes the postsynaptic response to the fusion of an individual SV with the AZ membrane. This parameter is reflected by the amplitude of spontaneously occurring miniature excitatory postsynaptic currents (minis) and can be influenced by the number and identity of postsynaptic receptors. We recorded similar mini frequencies and amplitudes at *wt* and *Cac^RNAi^* synapses (Figure 2R, S) demonstrating an unaltered quantal size and, in turn, suggesting a homeostatic addition of release sites. In line with a corresponding increase in postsynaptic receptor fields, puffing nicotine onto PNs elicited larger currents (Figure 2T).

Based on these results, we sought to match the functional increase of release sites in *Cac^RNAi^* to a structural correlate. To this end, we performed immunostainings of the antennal lobe and imaged Bruchpilot (Brp), a core component of the AZ cytomatrix (CAZ^50^), via confocal microscopy. To restrict our analysis to ORN-PN synapses, we co-labelled postsynaptic PNs via GFP and quantified Brp labels in the overlapping regions (Figure 2V). Consistent with the electrophysiological estimate of release site addition, the number of Brp clusters was significantly increased upon Cac knockdown (Figure 2U). Moreover, while the average cluster size remained unchanged in *Cac^RNAi^*, the signal intensity was significantly increased (Figure 2U), possibly reflecting the addition of CAZ units below the diffraction limit^51^. Thus, a homeostatic increase in the number of Brp-positive AZs counterbalanced the decreased transmitter release probability caused by reduced Cac expression.

### Monosynaptic effects and circuit activity underlie distinct features of coding reliability

Next, we examined whether the diminished coding reliability caused by *Cac^RNAi^* is due to monosynaptic mechanisms or instead arises from circuit effects. GABAergic iLNs also receive their major input from ORNs and inhibit these presynaptically. Whereas PNs are uniglomerular and receive homogenous ORN input, iLNs are multiglomerular and receive heterogeneous ORN input. PNs respond to ORN activity in a non-linear manner^30,43,52^. In contrast, GABAergic iLNs respond linearly to ORN activity^52^, are more sensitive to reduced input from ORNs^53^, and show a stronger transiency in their odor response^54^. Thus, we speculated that at least some of the observed reduction in PN reliability following presynaptic manipulations might arise from reduced recruitment of iLNs. In particular, this may explain the increased standard deviation in *Cac^RNAi^* that no longer occurred in the absence of the initial transient phase of PN odor responses (Figure 1R, S). To examine how iLN activity affects PN odor response reliability we compared *wt* and presynaptic *Cac^RNAi^* flies (Figure 3A). *Orco-GAL4* again drove *UAS-Cac^RNAi^* and *449-QF* was used to express *QUAS-GFP* in iLNs^55^. Reducing Cac protein levels in ORNs resulted in an almost complete abolishment of iLN responses to odors in both 0 and 2-4 days old flies (Figure 3B, C). Similar to PNs, *Cac^RNAi^* reduced the SV release probability at the ORN-iLN synapse (Figure 3D). However, in contrast to ORN-PN transmission, we observed no homeostatic rescue of iLN responses to odor stimulation even in 2-4 days old flies.

**Figure 3:**
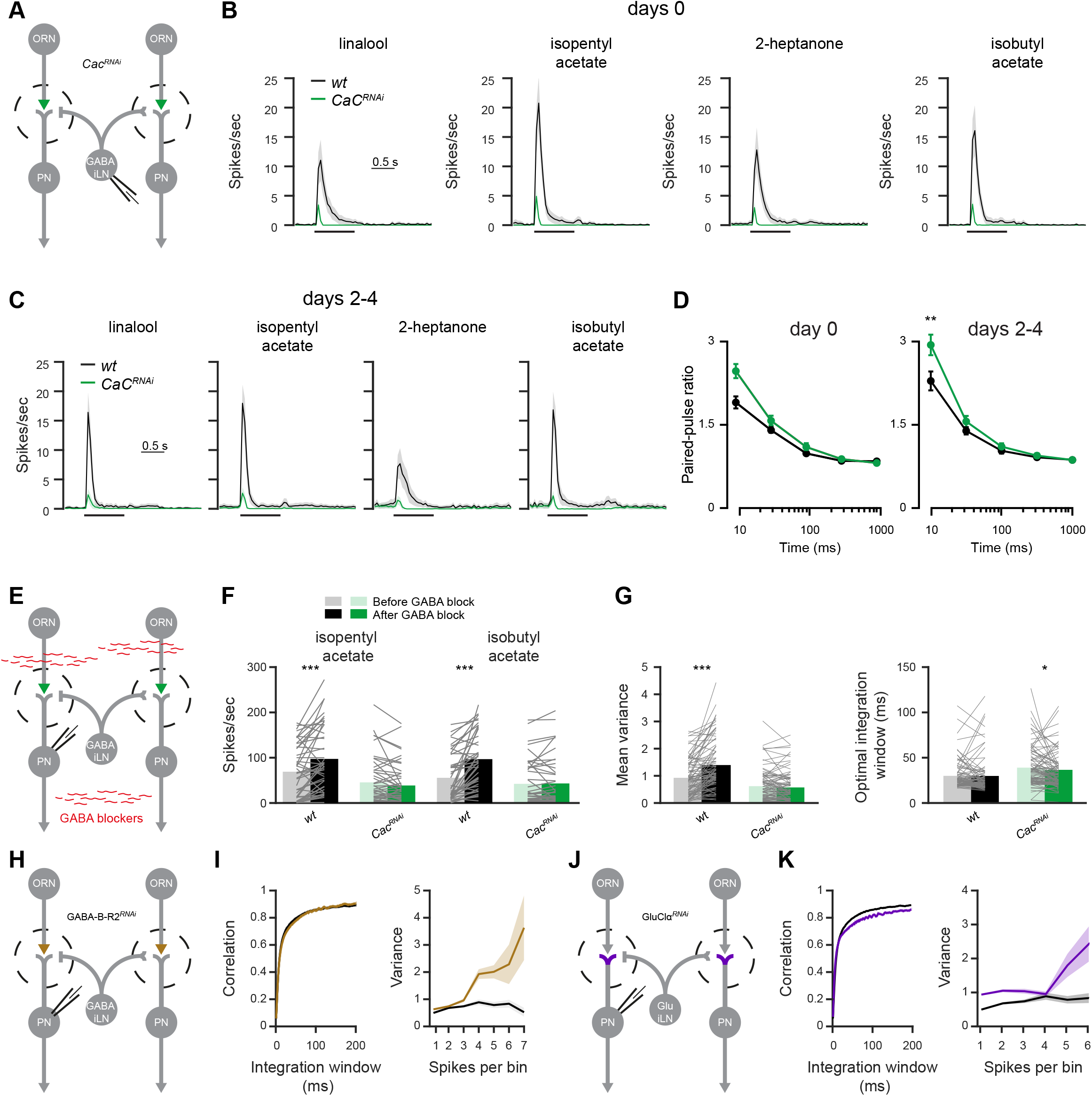
Monosynaptic effects underlie temporal reliability whereas circuit effects underlie rate code reliability. **A.** Experimental scheme. *Cac^RNAi^* was driven in ORNs using *Orco-GAL4*. Whole-cell patch recordings were made from iLNs labeled with the *449-QF* driver line. **B.** PSTH of the iLN population response to four odors examined as indicated (shaded areas represent SEM, the odor pulse is labeled with a black bar) for 0 day old flies. Cac knockdown resulted in an almost complete abolishment of iLN odor responses (final odor dilution of 5X10^−2^; n=49-50 flies). **C.** PSTH of the iLN population response to four odors examined as indicated (shaded areas represent SEM, the odor pulse is labeled with a black bar) for 2-4 days old flies. Cac knockdown resulted in an almost complete abolishment of iLN odor responses (final odor dilution of 5X10^−2^; n=49-50 flies). **D.** Quantification of the paired-pulse ratio at different inter-stimulus intervals for 0 day (left) and 2-4 days (right) old flies (10ms, 30ms, 100ms, 300ms and 1000ms). *Cac^RNAi^* significantly increased paired-pulse facilitation at short inter-pulse intervals (*wt*, n=13-22; *Cac^RNAi^*, n=15-28 flies). Error bars represent SEM. * p<0.05, two-sample t-test, see table S1. **E.** Experimental scheme. GABA receptors blockers (100 μM CGP54626 and 250 μM picrotoxin) were applied by bath perfusion. Whole-cell patch recordings were made from PNs labeled with the *GH146-QF* driver line. *UAS-Cac^RNAi^* was expressed in ORNs using *Orco-GAL4* **F.** Mean firing rate during the odor response. Application of GABA blockers significantly increased *wt* firing rates but had no effect on *Cac^RNAi^. Orco-GAL4* drove *UAS-Cac^RNAi^* and *GH146-QF* drove *QUAS-GFP* (n = 45-48 flies). **G.** *Left*. Temporal integration window and *right*, firing-rate reliability analysis for *wt* and *Cac^RNAi^* flies before and after the application of GABA receptors blockers (100 μM CGP54626, and 250 μM picrotoxin). The data were pooled for the presentation of two odors (isopentyl acetate and isobutyl acetate) and across all firing-rates (n=45-48 flies). **H.** Experimental scheme. An RNAi construct against the GABA-B-R2 receptor was expressed in ORNs using *Orco-GAL4*. Whole-cell patch recordings were made from PNs labeled with the *GH146-QF* driver line. **I.** *Left*. Temporal integration window and *right*, firing-rate reliability analysis for *wt* and *GABA-B- R2^RNAi^* flies (brown). The data were pooled for the presentation of two odors (isopentyl acetate and isobutyl acetate) and across all firing-rates (n=50-72 flies). **J.** Experimental scheme. An RNAi construct against the GluClα receptor was expressed in PNs using *GH146-GAL4*. Whole-cell patch recordings were made from PNs. **K.** *Left*. Temporal integration window and *right*, firing-rate reliability analysis for *wt* and GluClα^*RNAi*^ flies (purple). The data were pooled for the presentation of two odors (isopentyl acetate and isobutyl acetate) and across all firing-rates (n=50-72 flies). For all panels, * p<0.05, ** p<0.01, *** p<0.001, see table S1 for statistical analysis.

These results demonstrate that knocking down Cac in ORNs severely impairs the activation of iLNs. We therefore examined the effects of blocking GABAergic transmission on PN reliability. To this end, we exposed the brain to 100 μM CGP54626, a selective GABA_B_ receptor antagonist, and 250 μM picrotoxin, a chloride channel blocker^56^ and antagonist of GABA_A_ receptors (Figure 3E). As expected, GABA blockers affected *wt* firing rates (Figure 3F) but had no effect on PN spiking in *Cac^RNAi^* flies, where iLNs are hardly recruited (Figure 3B). When blocking GABA receptors in *wt* flies, we observed a significant increase in the variability of PN activity but the temporal integration window of PN responses was unaffected (Figure 3G). This suggests that iLNs affect only certain aspects of the code reliability. Since the odor response of iLNs was diminished when Cac was knocked down in ORNs we expected that blocking GABA receptors would not further affect coding reliability in *Cac^RNAi^* flies. Indeed, blocking GABAergic transmission had no effect whatsoever on variability and only a minor effect on the temporal integration window in presynaptic *Cac^RNAi^* (Figure 3G).

The AL also contains another population of inhibitory local neurons, which release glutamate. These glutamatergic interneurons mainly inhibit PNs, but were shown to have effectively similar roles as the GABAergic iLNs on the activity of the AL neural circuit^33^. Thus, the pharmacological experiments affect both the presynapse of ORNs and the postsynapse of PNs. Since iLNs mainly act by activating GABA-B receptors in the presynaptic terminal of ORNs^34,35^, we tested whether driving *GABA-B-R2^RNAi57^* in ORNs (Figure 3H) has similar effects on PN coding reliability as pharmacologically blocking the inhibitory iLNs (Figure 3G). Indeed, *GABA-B-R2^RNAi^* increased the variability at high firing rates but did not affect the temporal integration window (Figure 3I). The glutamatergic iLNs inhibit PNs by activating the glutamate receptor GluClα^33^. Therefore, we expressed an RNAi construct against GluClα in PNs (Figure 3J) and tested their odor response and coding reliability. Knockdown of the GluClα receptor in PNs had a strong effect on coding variability, as observed for the pharmacological experiments and by blocking the inhibitory presynaptic circuit (Figure 3K). In addition, we also observed an overall decrease in correlation (Figure 3K). However, in contrast to *Cac^RNAi^*, which increased PN integration time (Figure 1D), we did not observe any such increase in *GluClα^RNAi^*, if at all, the integration time of PNs was slightly decreased (Figure 3K and Figure S6). In summary, both pharmacological and genetic approaches suggest that the inhibitory AL circuit affects the variability of PN activity but not the temporal reliability.

### Reducing neural coding reliability disrupts behavioral reliability

The above results demonstrate that knocking down AZ Ca^2+^ channels specifically in ORNs decreases the coding reliability by PNs. To examine whether these changes are strong enough to affect behavioral output we tested the flies’ ability to correctly classify an ecologically relevant stimulus. To reduce response variability, which is often observed for naïve behavior^58,59^, we paired isopentyl acetate with an electric shock, using a well-established custom built apparatus^53,58–61^ and tested the accuracy of the animals’ behavioral responses, i.e. avoidance of isopentyl acetate. Following prolonged exposure (2 minutes) to the olfactory stimulus, presynaptic *Cac^RNAi^* animals were equally successful as their parental controls in classifying the odor (Figure 4A, B), indicating no impairment in the flies learning capabilities. However, when flies navigate in their natural habitat, olfactory stimuli are often brief and repetitive. Since such short stimuli could not be delivered in the above paradigm, we used an alternative assay in which the flies walked on a Styrofoam ball. This setting allowed us to present the same pattern of olfactory stimuli as used for the electrophysiological recordings (Figure 1). Ten one-second trials of the shock-paired odor isopentyl acetate were presented and the behavioral response was classified as correct if flies turned away from the odor source (Figure 4C). Indeed, *Cac^RNAi^* flies made significantly fewer correct choices as a result of identifying isopentyl acetate less reliably (Figure 4D). This reduction in correct stimulus classification was not due to a learning defect since the learning index was similar between *wt* and *Cac^RNAi^* flies (Figure 4B). Furthermore, Cac knockdown was only performed in the first order ORNs and not in third order neurons, which are required for memory formation^62^. Since our electrophysiological results showed that homeostatic synaptic plasticity rescues the neural code reliability only at lower odor concentrations, we tested whether behavioral reliability was also rescued at low stimulation intensities. The overall learning scores of *wt* and *Cac^RNAi^* flies were similar at low (Figure 4E) and high odor concentrations (Figure 4B). As predicted, *Cac^RNAi^* flies were as successful as *wt* animals in correctly classifying the low concentration odor (Figure 4F). In contrast, while *Cac^RNAi^* animals displayed normal memory formation at day 0 (Figure 4G), low concentration odor classification was significantly impaired in young flies (Figure 4H). This nicely matches our finding that homeostatic compensation takes a day to develop. In summary, these results demonstrate how synaptic plasticity can maintain neural coding reliability and enable the animal to consistently classify an important physiological stimulus.

**Figure 4:**
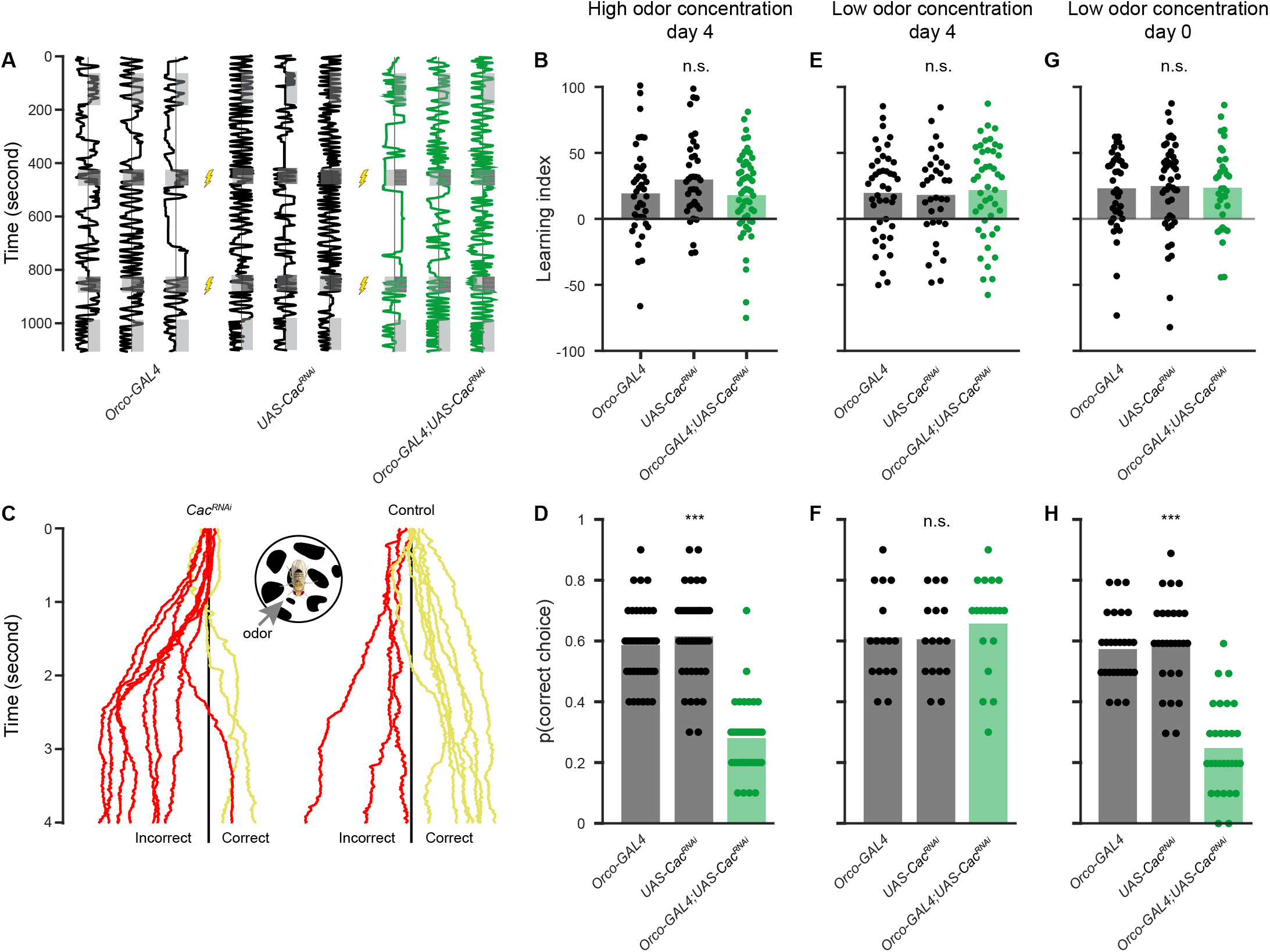
Cac knockdown reduces behavioral reliability. **A.** Experimental scheme for the learning paradigm. Flies were constrained in a linear chamber with isopentyl acetate presented on one side (gray). For pairing with an electric shock, the odor was presented on both sides of the chamber. Examples of single fly trajectories are shown. **B.** Learning performance for prolonged odor exposure at high odor concentration. No significant differences in the learning index were observed between the parental controls and *Cac^RNAi^* in ORNs. Each dot represents a single fly (final odor dilution of 5X10^−2^; n=37-54). **C.** Trained flies were tested in an assay where flies walked on a ball and the shock-paired odor was presented from the indicated side. Ten one second odor trials were presented as in Figure 1. Examples for single fly trajectories along the virtual X-axis are presented. Choice was classified as correct (yellow) if the mean X-axis position following the odor presentation was rightward of the X-axis midline (black line) and incorrect (red) otherwise. **D.** Behavioral reliability measure. The behavioral reliability measure was defined as the probability of correctly classifying the shock-paired odor. Cac knockdown in ORNs reduces the behavioral reliability compared to the parental controls at high odor concentration. Each dot represents a single fly (final odor dilution of 5X10^−2^; n=20-21). **E.** Learning performance for prolonged odor exposure at low odor concentration in 4 days old fly. No significant differences in the learning index were observed between the parental controls and *Cac^RNAi^* in ORNs. Each dot represents a single fly (final odor dilution of 5X10^−4^; n=34-45). **F.** Behavioral reliability measure as in panel D for low odor concentration in 4 days old fly. Cac knockdown in ORNs did not affect the behavioral reliability compared to the parental controls at low odor concentration. Each dot represents a single fly (final odor dilution of 5X10^−4^; n=16-19). **G.** Learning performance for prolonged odor exposure at low odor concentration in 0 day old flies. No significant differences in the learning index were observed between the parental controls and *Cac^RNAi^* in ORNs. Each dot represents a single fly (final odor dilution of 5X10^−4^; n=34-47). **H.** Behavioral reliability measure for low odor concentration in 0 day old flies. Cac knockdown in ORNs reduces the behavioral reliability compared to the parental controls at low odor concentration, indicating a lack of homeostatic compensation. Each dot represents a single fly (final odor dilution of 5X10^−4^; n=27-29). For all panels *** p<0.001, see table S1 for statistical analysis.

## Discussion

In this study, we addressed two distinct yet related questions. First, we provide *in vivo* evidence that the molecular integrity of the AZ is important for neural coding reliability. Second, we demonstrate that homeostatic synaptic plasticity operates at the ORN-PN synapse and that this process compensates a reduction in release probability by increasing the number of ORN AZs. The homeostatic adjustment restored eEPSC amplitudes in PNs and neural coding reliability. We further show that the restored neural reliability maintains behavioral performance. Finally, we demonstrate that the homeostatic compensation is limited in its efficiency and fails to maintain reliable neural coding and behaviour at elevated firing frequencies in response to high odor concentrations.

Using *in vivo* recordings, we show that high release probability of ORN AZs is required for a reliable neural code in the postsynaptic PNs in response to physiological odor stimuli. Knockdown of Cacophony from ORN AZs that reduced ORN release probability affected neural code reliability in several manners. First, the high firing rates of the initial transient phase of PN odor responses became more variable due to reduced recruitment of iLNs. Second, the onset of odor responses was delayed and more variable. Third, the temporal dynamics of the odor response became less reliable. This third effect results most probably from monosynaptic changes to ORN-PN transmission. At a behavioral level, we find that reduced coding reliability of PNs impairs the animals’ ability to correctly classify a physiologically relevant stimulus.

We found that knocking down Cacophony affected neuronal coding reliability mainly at high firing rates. This was true for both the monosynaptic as well as the circuit mechanisms. In terms of the circuit effects on rate code reliability, this is in line with the recruitment of iLNs. iLNs sample many types of ORNs with different response profiles^48^ and as a result, they respond linearly to ORN activity^52^. Thus, iLNs respond most strongly to high ORN firing rates, which in turn are triggered by high odor intensities. Correspondingly, almost entirely eliminating the recruitment of iLNs by manipulating ORN AZs (Figure 3), mostly affects processing of high firing rates. We find that a decrease in SV release probability increases the latency and jitter of eEPSC (Figure 2), accompanied by an increase in the latency and jitter of postsynaptic action potential initiation (Figure 1). We further show that this drop in the temporal precision of synaptic transmission increases in a frequency-dependent manner. This finding helps to explain why disrupted AZ function mainly impairs temporal coding at high firing rates and strong odor intensities.

Theoretical studies have suggested that decreasing SV release probability should result in reduced neural coding reliability^16–20,63–65^, although exceptions have also been demonstrated^45,66^. However, whether these theoretical studies hold true *in vivo*, with highly dynamic synaptic properties, in complex neural circuits, and in response to physiological stimuli has remained an open question. To the best of our knowledge, the present study provides the first *in vivo* evidence that interfering with the molecular control of SV release results in reduced neural coding reliability. Notably, the ORN-PN synapse is surprisingly resilient to reduced AZ Ca^2+^ channel expression and compensates the drop in release probability by increasing the number of AZs to yield normal eEPSC amplitudes. Homeostatic synaptic plasticity features at the *Drosophila* NMJ^11,12^, mushroom body^67^, and also operates in the antennal lobe to match synaptic strength to PN excitability^30,68^. Interestingly, *Cac^RNAi^* decreases synaptic transmission to a greater extent from ORNs to iLNs than from ORNs to PNs (compare Figures 1C and 3C). Thus, the functional impact of Cac or mechanisms to compensate for reduced Ca^2+^ channel expression differ between ORN AZs depending on the identity of the postsynaptic partner. Interestingly, a recent study demonstrated that during the time window of homeostatic compensation, i.e. the first day post-eclosion, ORNs and PNs show structural changes involving increased Brp expression and neurite expansion, while iLNs do not^69^. It is conceivable that this developmental stability prevents or limits structural homeostatic plasticity at ORN-iLN synapses. Moreover, parallels may exist with the NMJ, where the expression of presynaptic homeostatic plasticity is compartmentalized to a subset of a motoneuron’s synapses depending on their target muscle^70^. These considerations emphasize the importance of further future studies on the molecular heterogeneity of AZs in the olfactory system^4,71,72^ and of elucidating the physiological properties of individual synapses in the context of neural information processing.

## Methods

### Fly Strains

Fly strains (see below) were raised on cornmeal agar under a 12 h light/12 h dark cycle at 25 °C. The following fly strains were used: *UAS-Cac^RNAi^* ^73^ (VDRC ID 5551), UAS-GluClα^RNAi^ (53356), UAS-GABA-B-R2^RNAi^ (50608), *GH146-QF,QUAS-mCD8-GFP* (BDSC_30038), *Orco-GAL4* (BDSC_26818), *UAS-GCaMP6f* (BDSC_52869), *449-QF* ^55^, *QUAS-mCD8-GFP* (BDSC_30002)

### Olfactory stimulation

Odors (purest level available) were obtained from Sigma-Aldrich (Rehovot, Israel). Odor flow of 0.4 l/min (10^−1^ dilution) was combined with a carrier air stream of 0.4 l/min using mass-flow controllers (Sensirion) and software controlled solenoid valves (The Lee Company). This resulted in a final odor dilution of 5X10^−2^ delivered to the fly. Odor flow was delivered through a 1/16 inch ultra-chemical-resistant Versilon PVC tubing (Saint-Gobain, NJ, USA) placed 5 mm from the fly’s antenna.

### Electrophysiology

Flies were anesthetized for 1 minute on ice. A single fly was fixed to aluminum foil using wax. To expose the brain, the cuticle and trachea were removed. The brain was superfused with carbonated solution (95% O_2_, 5% CO_2_) containing 103 mM NaCl, 3 mM KCl, 5 mM trehalose, 10 mM glucose, 26 mM NaHCO_3_, 1 mM NaH_2_PO_4_, 1.5 mM CaCl_2_, 4 mM MgCl_2_, 5 mM N-Tris (TES), pH 7.3. For *in vivo* whole-cell recordings 0,1 or 2-4 days old flies were used as previously described ^74^. Briefly, flies’ brains were visualized on a Scientifica SliceScope Pro 1000 upright microscope with a 40x water immersion objective. Patch pipettes of 9–12 MΩ resistance were used. Intracellular solution contained: potassium aspartate 140mM, HEPES 10mM, KCL 1mM, MgATP 4mM, Na_3_GTP 0.5mM, EGTA 1mM with pH of 7.3 and osmolarity of 265 mOsm. PNs were randomly patched from both the lateral and medial cluster and iLNs were randomly patched from the lateral cluster. Voltage or current recordings were performed using Axon Instruments MultiClamp 700B amplifier in current- or voltage-clamp mode respectively. Data was low-pass filtered at 1 kHz and sampled at 50 kHz. Upon break-in to the cell, a small constant current was applied to maintain a membrane potential of −60mV. Action potential times were extracted using a custom MATLAB code followed by manual inspection to verify correct identification. Spike times were aligned to the beginning of the rising phase of the membrane potential, indicating the beginning of the odor stimulus. In experiments where GABA blockers were used, 100 μM of CGP 54626 (Tocris, CAS: 149184-21-4) and 250 μM of picrotoxin (Sigma-Aldrich, CAS: 124-87-8) were bath applied. eEPSCs were evoked by stimulating ORN axons with a minimal stimulation protocol via a suction electrode ^30^. Brief (typically 50μs) pulses were passed through the innervating nerve using a constant current stimulator (Digitimer, DS3 Isolated Current Stimulator). To measure eEPSC amplitude and kinetics 32 eEPSCs were evoked at 1 Hz and averaged. eEPSC latency was measured from the beginning of the stimulation artifact to the beginning of the eEPSC rising phase. Paired-pulse recordings were made at 0.2 Hz with inter-stimulus intervals of (in ms): 10, 30, 100, 300 and 1,000. For each interval 20 traces were averaged. Ten seconds of rest were afforded to the cell in between recordings. The amplitude of the second response in 10 ms inter-pulse recordings was measured from the peak to the point of interception with the extrapolated first response. Data were analyzed using MATLAB.

*In vivo* patch-clamp recordings of minis were performed essentially as previously described^75^. Cell bodies of PNs were visualized with a Scientifica Slice Scope equipped with a 40 x Water Objective (Olympus 40x, NA 0.8). Electrodes (6 - 8MOhm) were filled with internal solution containing (in mM): Potassium aspartate 125, CaCl_2_ 0.1, EGTA 1.1, HEPES 10, MgATP 4, Na_3_GTP 0.5, pH adjusted to 7.3, osmolarity was 265 mOsm. To visualize PN morphology, Biocytin (3 mg/ml) was added to the intracellular solution. Throughout measurements, preparations were perfused with oxygenated external saline, containing (in mM): NaCl 103, KCl 3, CaCl_2_ 1.5, MgCl_2_ 4, NaH_2_PO_4_ 1, NaHCO_3_ 26, TES 5, Trehalose 10, Glucose 10, 277 ± 2 mOsm, pH 7.3. Additionally, 4 μM TTX (Carl Roth, 6973) was supplied to the bath solution to suppress spontaneous spiking. For recordings, PNs of female flies were held at a command potential of - 80 to - 100 mV. Signals were low-pass filtered at 1kHz and digitized at 10kHz. For detection of minis, templates were generated and applied to recordings (ClampFit 11.0.3).

### Functional Imaging

Flies used for functional imaging were reared as described above. Imaging was done as previously described ^59,61,76^. Briefly, using two-photon laser-scanning microscopy (DF-Scope installed on an Olympus BX51WI microscope). Flies were anesthetized on ice then a single fly was moved to a custom built chamber and fixed to aluminum foil using wax. Cuticle and trachea in the required area were removed, and the exposed brain was superfused with carbonated solution as described above. Odors at final dilution of 5X10^−2^ were delivered as described above. Fluorescence was excited by a Ti-Sapphire laser (Mai Tai HP DS, 100 fs pulses) centered at 910 nm, attenuated by a Pockels cell (Conoptics) and coupled to a galvo-resonant scanner. Excitation light was focused by a 20X, 1.0 NA objective (Olympus XLUMPLFLN20XW), and emitted photons were detected by GaAsP photomultiplier tubes (Hamamatsu Photonics, H10770PA-40SEL), whose currents were amplified (Hamamatsu HC-130-INV) and transferred to the imaging computer (MScan 2.3.01). All imaging experiments were acquired at 30 Hz.

### Immunohistochemistry

Whole mount stainings were performed essentially as previously described^77^. For analyzing Brp, brains were fixed with 4% PFA for 2h, washed with PBT (0.3%) and blocked using 5% normal goat serum in PBT overnight at 4°C, before mouse-anti Brp [nc82, 1:50; provided by E. Buchner; RRID: AB_528108^78^ was added. Following another overnight incubation at 4°C, samples were washed with PBT before goat anti-mouse STAR RED (1:200; Abberior #2-0002-011-2, RRID:AB_2810982) was applied overnight. After washes with PBT, samples were stored in mounting medium (Abberior MOUNT, LIQUID ANTIFADE, #MM-2009). To preserve tissue morphology, imaging spacer (Sigma-Aldrich #GBL654008) were used. For verification of Cacophony knock down at ORN pre-synapses, we followed a protocol published by Chang and colleagues^79^. Whole mounts were fixed with Bouins fixative (Carl Roth #6482.3), washed with PBS and blocked in PBT (0.2%) with 5% NGS at 4°C overnight. This was followed by an overnight incubation with rabbit anti-cacophony (1:1000) in blocking solution, before the samples were washed several times with PBT. Subsequently, whole mounts were incubated with biotinylated goat anti rabbit IgG (1:500; Jackson ImmunoResearch Labs #111-065-003, RRID:AB_2337959), washed with PBT and subsequently incubated with STAR RED streptavidin (1:500; Abberior #STRED-0120). Samples were stored in mounting medium (Abberior MOUNT, LIQUID ANTIFADE, #MM-2009) until final use.

### Confocal microscopy

Images were acquired with an Abberior INFINITY LINE system (upright Olympus BX63F) equipped with a 60x / NA 1.42 oil immersion objective. Laser settings were kept constant and image acquisition alternated between genotypes. Image analysis was done using ImageJ (National Institutes of Health, Bethesda). Analysis was performed on each image in a stack. Data sets exceeding the maximal slice number of 125 slices per stack were excluded from analysis. To restrict image analysis to signals at ORN-PN synapses, masks of the GFP signal (driven with GH146) were created and overlayed with the Brp or Cacophony channel. Individual punctae were detected with the “Find Maxima” command and quantified via “Analyze Particles”.

### Behavioral chambers

Experiments were performed using a custom-built, fully automated apparatus ^58,60,61^. Single flies were housed in clear chambers (polycarbonate, length 50 mm, width 5 mm, height 1.3 mm). Mass flow controllers (CMOSens PerformanceLine, Sensirion) were used to control air flow. An odor stream (0.3 l/min) obtained by circulating the air flow through vials filled with a liquid odorant was combined with a carrier flow (2.7 l/min). Isopentyl acetate was prepared at 10 fold dilution in mineral oil. Liquid dilution and mixing carrier and odor stimulus stream resulted in a final 100 fold dilution. Fresh odors were prepared daily.

Two identical odor delivery systems were used each delivering odors independently to each half of the chamber. The total flow (3 l/minute, carrier and odor stimulus) was split between 20 chambers.. The air flow from the two halves of the chamber converged at a central choice zone. The 20 chambers were stacked in two columns each containing 10 chambers and were backlit by 940 nm LEDs (Vishay TSAL6400). Images were obtained by a MAKO CMOS camera (Allied Vision Technologies) equipped with a Computar M0814-MP2 lens. The apparatus was operated in a temperature controlled incubator (Panasonic MIR 154) at 25°C.

Fly position was extracted from video images using a virtual instrument written in LabVIEW 7.1 (National Instruments). The same virtual instrument was also used to control odor delivery. Data were analyzed in MATLAB 2018a (The MathWorks). Conditioning protocol included 12 equally spaced 1.25 s electric shocks at 50 V, and was repeated twice. The learning index was calculated as (preference for isopentyl acetate before training) – (preference for isopentyl acetate after training). Flies that were used for the ball assay in figure 4C were given a training protocol that consisted of 12 equally spaced 1.25 s electric shocks at 50 V, and repeated 6 times with 15 minutes interval. Flies were then tested on the ball assay 12 hours later.

### Ball assay

Flies walked on a treadmill ball as previously described ^80^. Briefly, the fly thorax was glued to a fine metal rod using wax in a manner that allowed for free limb movement. Flies were then placed on a Styrofoam ball with a diameter of 9 mm which floated on an air-steam in a custom made ball holder. Videos were captured using a Blackfly S (FLIR^®^ Systems) camera fitted with a Computar Macro zoom 0.3–1×, 1:4.5 lens at a frame rate of 100 FPS. The ball was illuminated by infrared LEDs and the fly position was tracked offline using FicTrac ^81^. Odor delivery was the same as for the electrophysiological recording experiments described above.

## Quantification and statistical analysis

### Integration wlndow analysis

For this analysis, only the first two seconds after the beginning of the odor stimulus were used. Spike trains were then binned with bin sizes ranging from 1ms to 200ms with 1ms intervals. For each bin size, the pairwise Pearson correlation was calculated for all combinations of the 10 trials for each odor. Trials that did not show any odor response were excluded from this analysis. Integration window was defined by using the Youden’s index ^82^, which is the point of maximal distance between the correlation vs. bin size curve and the diagonal between the minimum and maximum point of said curve. The Youden’s index is a communally used measure for the optimal cutoff point of a curve, such as the one obtained by plotting correlation vs. bin size^83^.

### Variance analysis

For each fly, the mean and variance of the 10 stimuli repetitions were calculated using time bins of 20ms. The mean spike count of each bin was then binned into discrete bins along with the corresponding variance in that time bin.

### Statistics and data analysis

All statistical testing and parameter extraction were done using custom MATLAB code (The MathWorks, Inc.). All statistical tests details can be found in table S1. Significance was defined as a p-value smaller than 0.05 and all statistical tests were two-sided. Normality assumption was tested using the Shapiro-Wilk test (https://www.mathworks.com/matlabcentral/fileexchange/13964-shapiro-wilk-and-shapiro-francia-normality-tests). In cases where the normality assumption was violated a permutation test was used using the ‘permutationTest’ function in MATLAB (https://github.com/lrkrol/permutationTest).

Effect size was calculated with the Measures of Effect Size (MES) Toolbox https://github.com/hhentschke/measures-of-effect-size-toolbox/blob/master/readme.md). Permutation test was used using the ‘permutationTest’ function in MATLAB (https://github.com/lrkrol/permutationTest).

For presentation, bar plots with dots were generated using the UnivarScatter MATLAB ToolBox (https://www.mathworks.com/matlabcentral/fileexchange/54243-univarscatter), raster plots were generated with the Flexible and Fast Spike Raster Plotting ToolBox (https://www.mathworks.com/matlabcentral/fileexchange/45671-flexible-and-fast-spike-raster-plotting) and the shadedErrorBar function (https://github.com/raacampbell/shadedErrorBar) for shaded errors on imaging traces.

## Data and code availability

The data and code used to generate the figures in this manuscript are available upon publication in the first author GitHub page: https://github.com/Eyal-ro.

## Acknowledgments

We thank Dr. Ya-Hui Chou and the Bloomington Stock Center for fly strains. We also thank David Morton for kindly sharing the Cacophony antibody. This work was supported by the European Research Council (676844, MP) and the Deutsche Forschungsgemeinschaft (project numbers 430156010/SPP 2205 to RJK and 408264519 to MP and RJK).

## Author contributions

ER: conceptualization, methodology, investigation, formal analysis, software, writing–review & editing, visualization. NE: investigation, formal analysis, writing–review & editing, visualization JEM: investigation, formal analysis, visualization. RJK: Initiated the project, conceptualization, methodology, writing–original draft, writing–review & editing, supervision, funding acquisition. MP: Initiated the project, conceptualization, methodology, investigation, formal analysis, software, writing–original draft, writing–review & editing, visualization, supervision, funding acquisition.

## Supplementary Information

**Figure S1:**
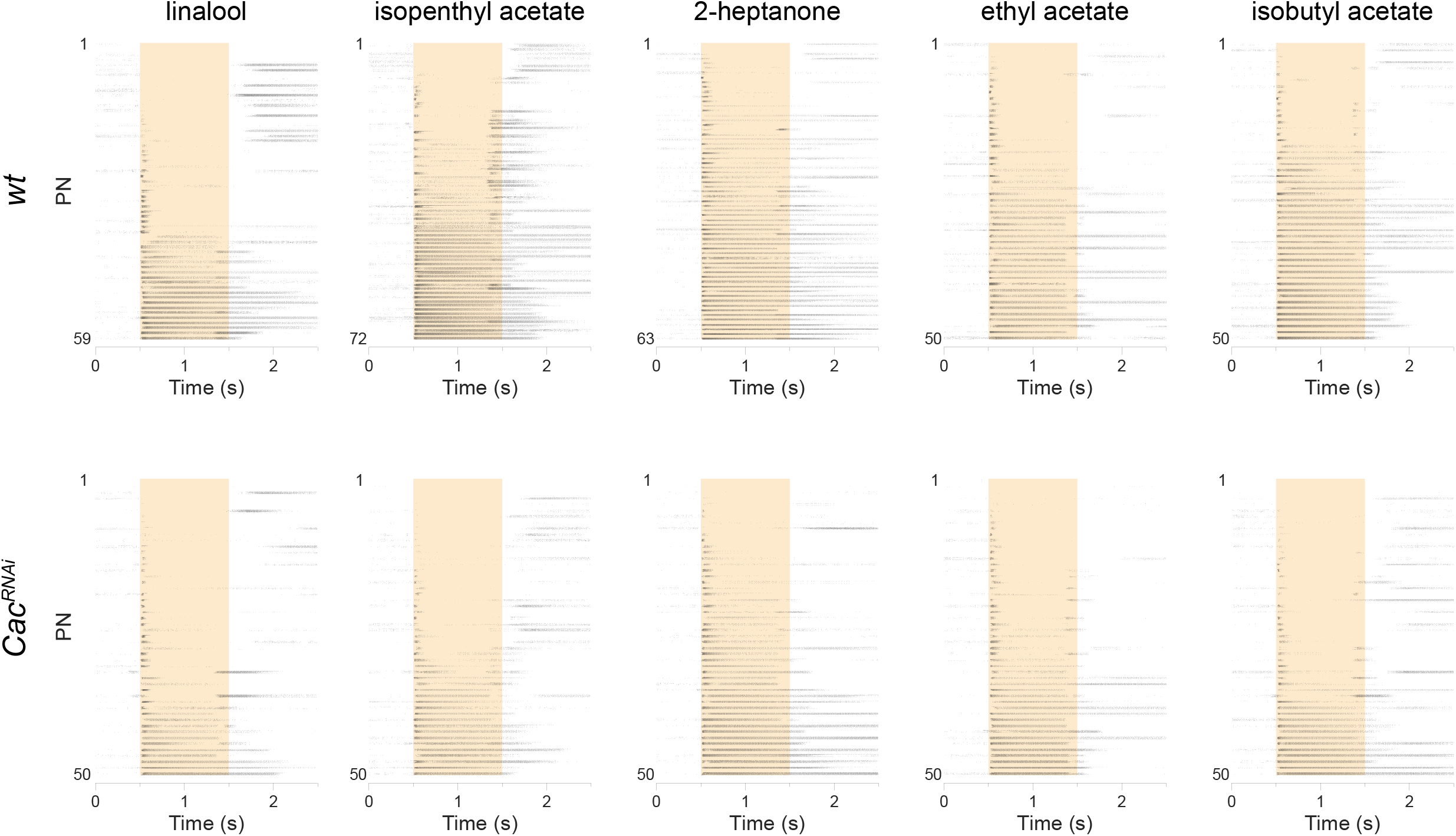
PN population responses to odor stimulation. Raster plots of PN population responses to five odors as indicated (final odor dilution of 5X10^−2^) in *wt* flies and *UAS-Cac^RNAi^* in ORNs. Each neuron was presented with 10 repetitions of the olfactory stimulus. *Orco-GAL4 UAS-Cac^RNAi^* in ORNs and *GH146-QF* drove *QUAS-GFP* (n=45- 72).

**Figure S2:**
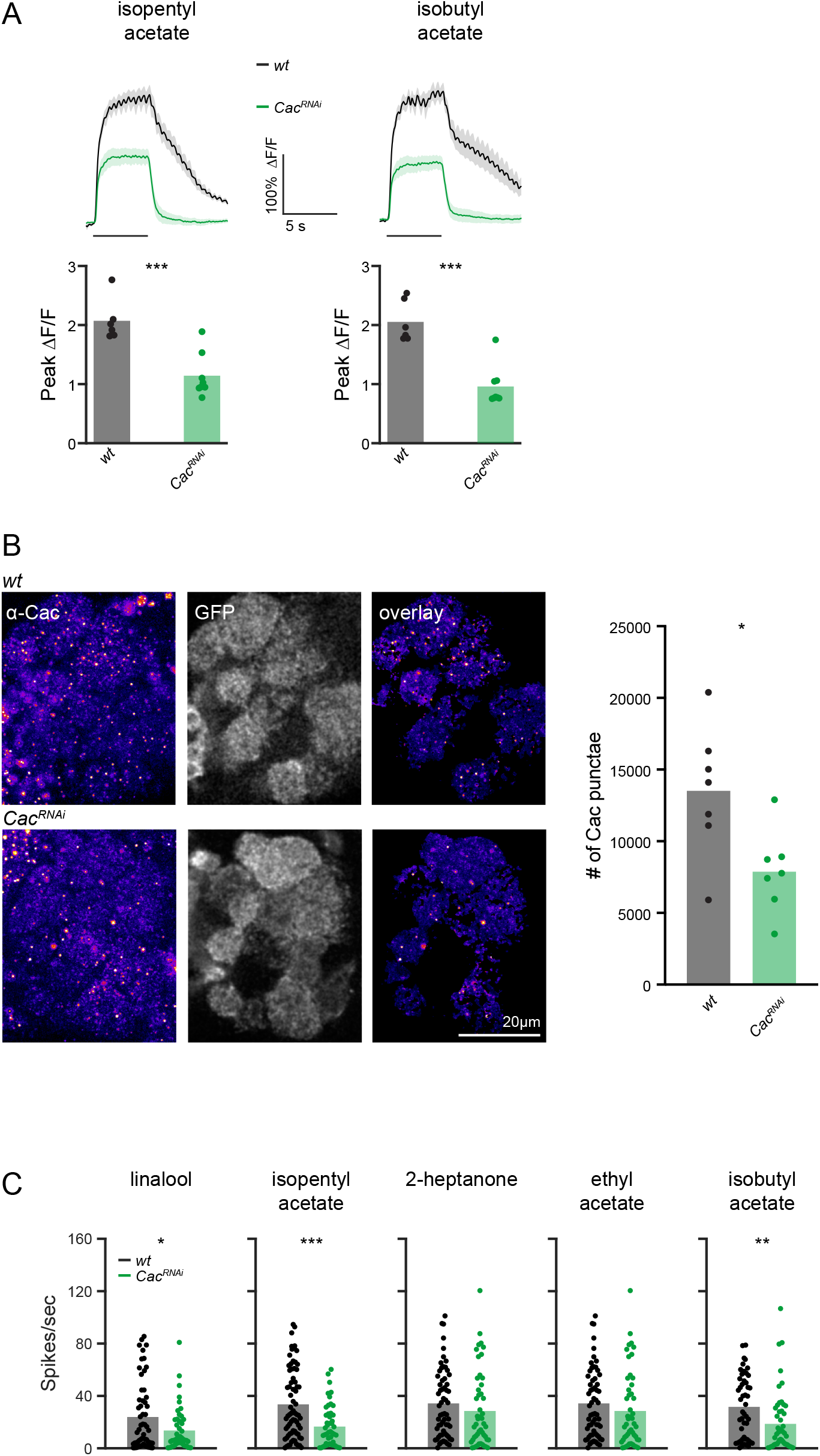
Knockdown of Cac affects ORNs calcium levels and reduces odor-stimulated PN firing rates. **A.** *Top*, averaged traces ± SEM (shading) of odor responses (as indicated, odor pulse is labeled with a black bar) obtained from a single plane of the entire AL for *wt* or RNAi flies. *Orco-GAL4* drove *UAS-Cac^RNAi^* along with *UAS-GCaMP6f. Bottom*, peak ΔF/F during odor responses for the traces presented in the *top* panel. As expected, a significant decrease in Ca^2+^ signals was observed for *Cac^RNAi^*. (n = 5-7 flies, ** p < 0.01, ** p < 0.001, Two-sample t-test, see table S1). **B.** To validate knock-down of Cacophony at ORN-PN synapses, whole mounts of flies were stained with an antibody against Cacophony^1^ (Chang et al., 2014) in control (upper panel; *orco- Gal4/+; GH-146-QF,QUAS-GFP/+)* and Cac^RNAi^ flies (lower panel*; orco-Gal4/+; GH146- QF,QUAS-GFP/UAS-Cac^RNAi^)*. Analysis of Cac signals (fire) was restricted to excitatory PN post-synapses trough an overlay with the respective mask generated by imaging GH146 driven GFP (grey, GFP; overlay). Scale bar 20μm. *Right*. Confocal analysis yielded a lower number of Cac signals at ORN-PN synapses upon Cac knock-down in ORNs, confirming the RNAi efficacy. (n=7, * P ≤ 0.05). **C.** Mean firing rate of PNs in response to five odors as indicated (final odor dilution of 5X10^−2^) for *wt* flies and for *Cac^RNAi^*. Each data point shows the average of 10 responses to a 1 s odor stimulus and 1 s following the odor stimulus. *Orco-GAL4* drove *UAS-Cac^RNAi^* in ORNs and *GH146-QF* drove *QUAS-GFP*. For most odors a significant decrease in firing rate was observed. (n=45-72, * p<0.05, ** p<0.01, *** p<0.001, Two-sample t-test, see table S1).

**Figure S3:**
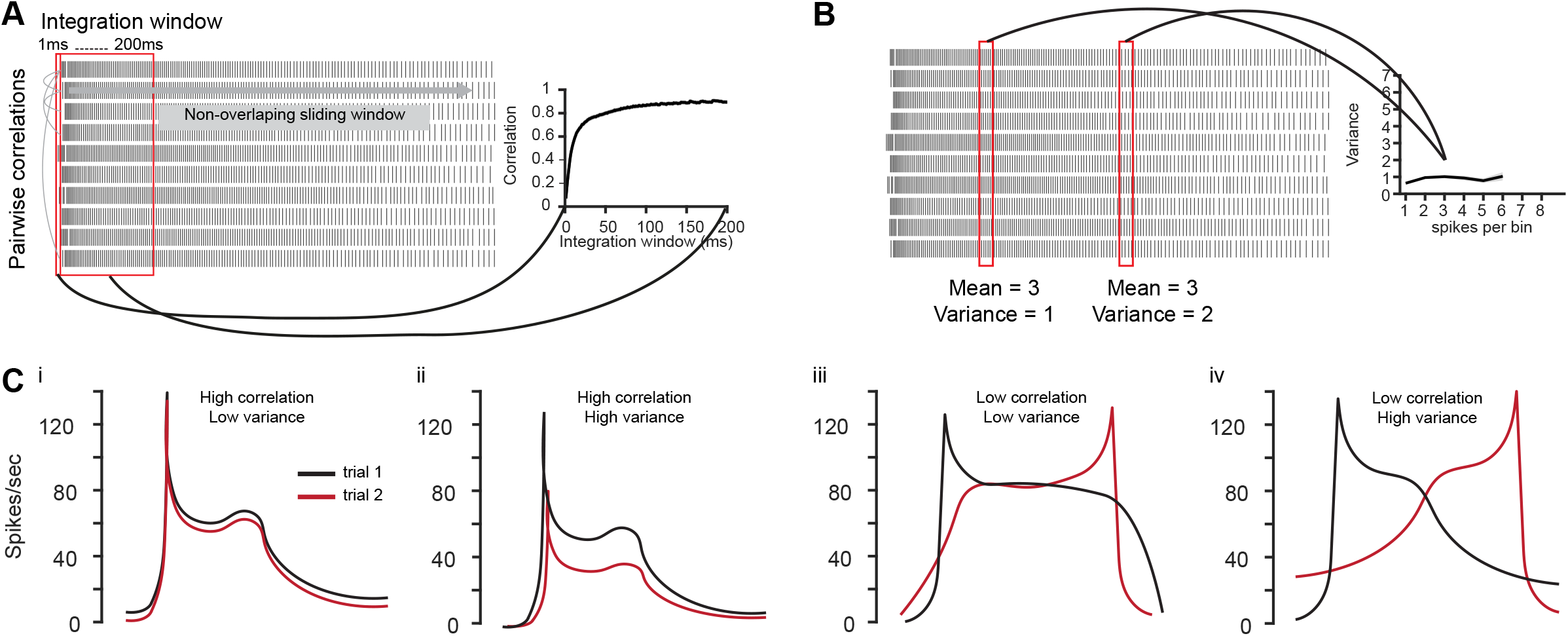
Graphical description of the reliability analysis. **A.** Temporal reliability analysis. Pairwise correlations were performed between the 10 repetitions of each odor-neuron combination. Correlations were calculated using increasing, non-overlapping integration windows from 1 to 200 ms with 1 ms intervals. The 45 pairwise correlation values for each condition (i.e. PN and odor) were averaged to a single correlation value for this condition. All correlation values for all PNs and odors were pooled to yield the correlation curve on the right. **B.** Firing rate reliability analysis. Spike trains were binned using 20 ms windows. The number of spikes and variability was calculated for each individual bin. The mean variance value for all bins having the same number of spikes is presented on the variance curve on the right. Variance values are pooled across neurons, odors and time. **C.** Temporal reliability analysis and firing rate reliability analysis capture different aspects of trial-to-trial coding reliability. Correlation captures the general shape of the PSTH regardless of momentary changes in firing rate at a given point in time (i and ii). Inter trial variability captures the momentary changes in firing rates regardless of the PSTH temporal dynamics (ii and iv).

**Figure S4:**
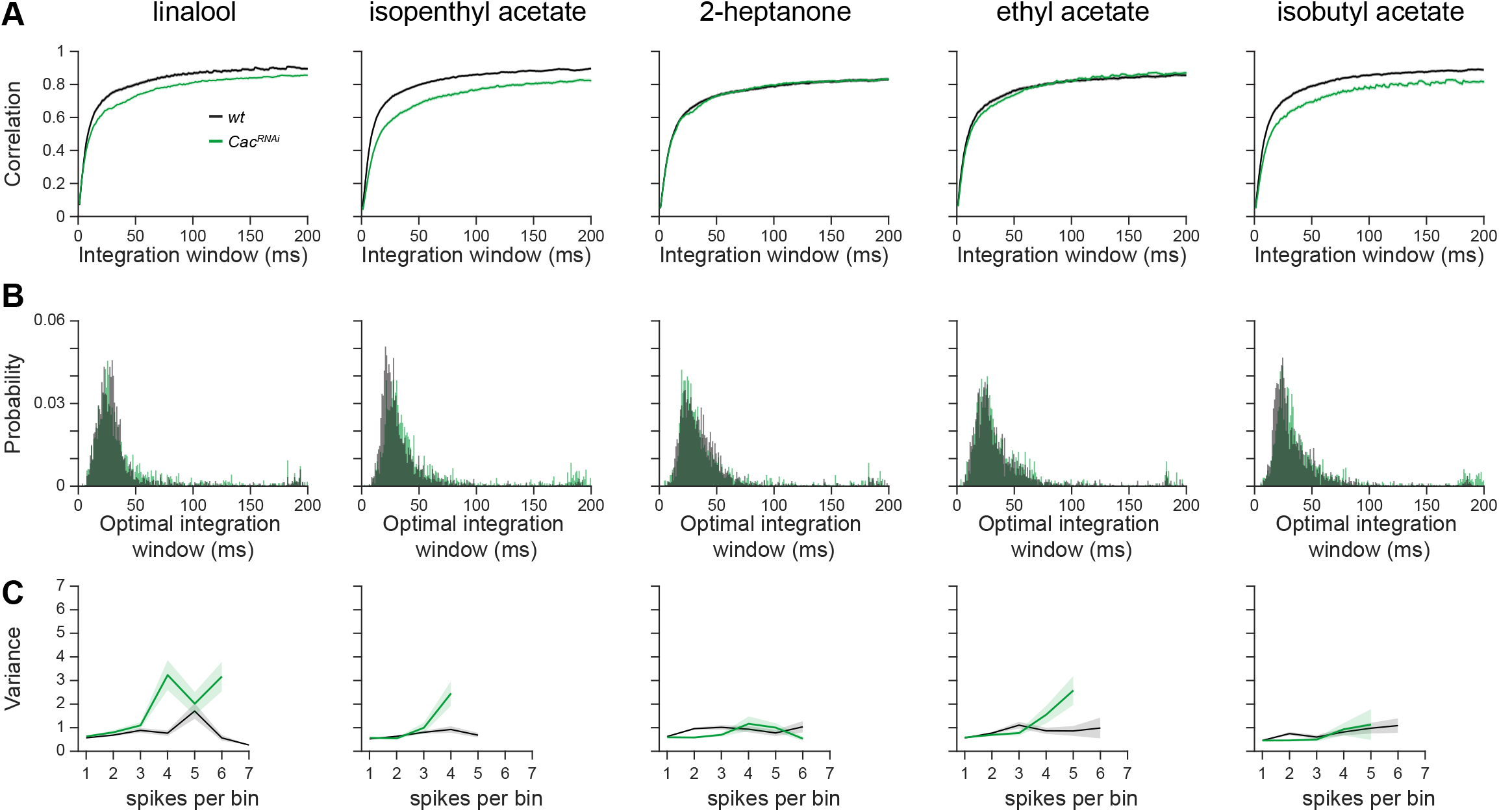
Single odor analyses of response reliability. **A.** Temporal reliability analysis as performed in Figure 1A but for single odors as indicated. Data obtained from Figure 1 and Figure S1. For all odors, a final odor dilution of 5X10^−2^ was used. **B.** The optimal integration window analysis as performed in Figure 1D but for single odors as indicated. Data obtained from Figure 1 and Figure S1. For all odors, a final odor dilution of 5X10^−2^ was used. **C.** Firing rate reliability analysis as performed in Figure 1E but for single odors as indicated. Data obtained from Figure 1 and Figure S1. For all odors, a final odor dilution of 5X10^−2^ was used.

**Figure S5:**
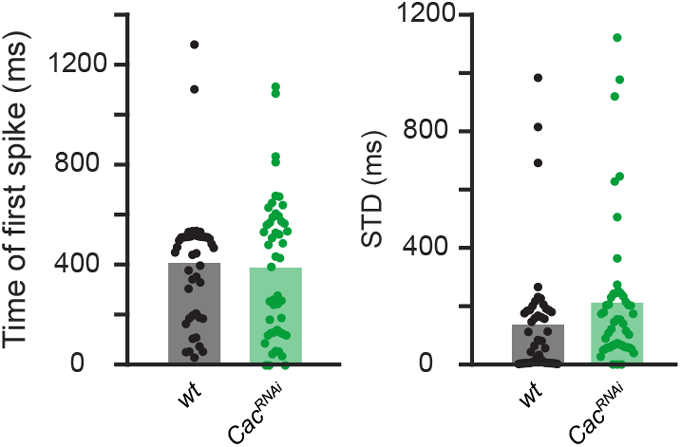
Controls for first spike analysis. Analysis of the latency of the first spike (left) and jitter of the first spike (right) in PNs as performed in Figure 1I,J but in response to mock odor application (i.e. no odor was actually applied) for *wt* flies or *Cac^RNAi^* in ORNs (n=47-49). No change in any parameter was observed.

**Figure S6:**
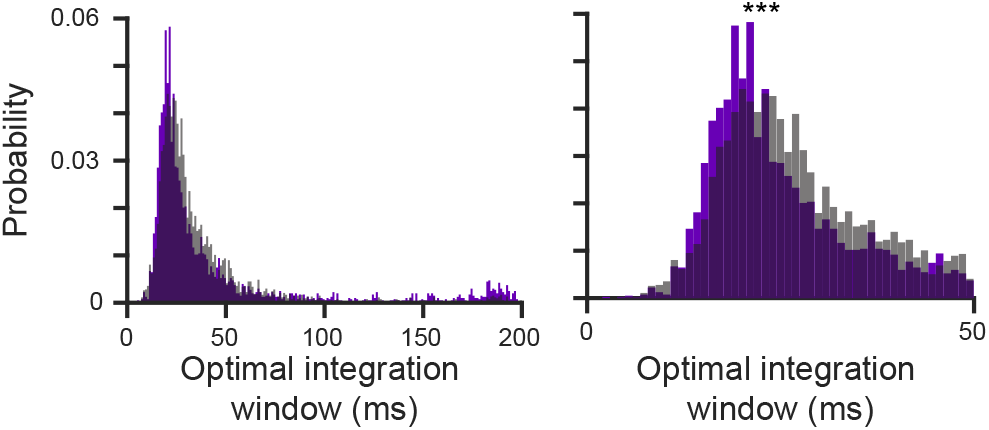
GluClα RNAi in PNs does not increase their integration window. *Left*, the curve saturation point was calculated for each odor-neuron combination and pooled across the two odors for the data in Figure 3K. *Right*, Left curves are presented at a larger scale. GluClα knockdown in PNs did not increase their optimal temporal integration window. If at all, it slightly improved as reflected by a smaller integration window spread and a slight peak shift to the left.

---

**Table S1:**
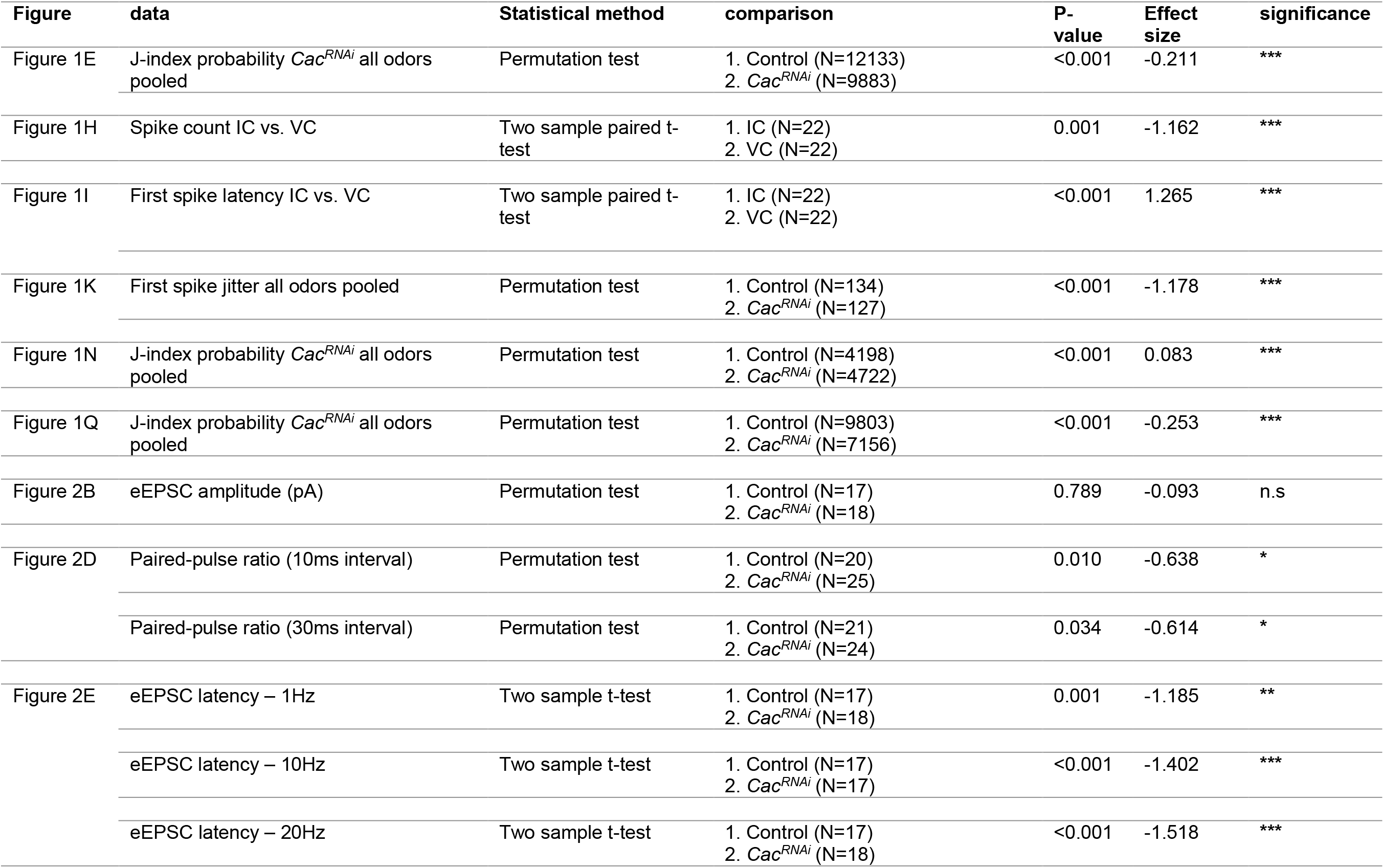
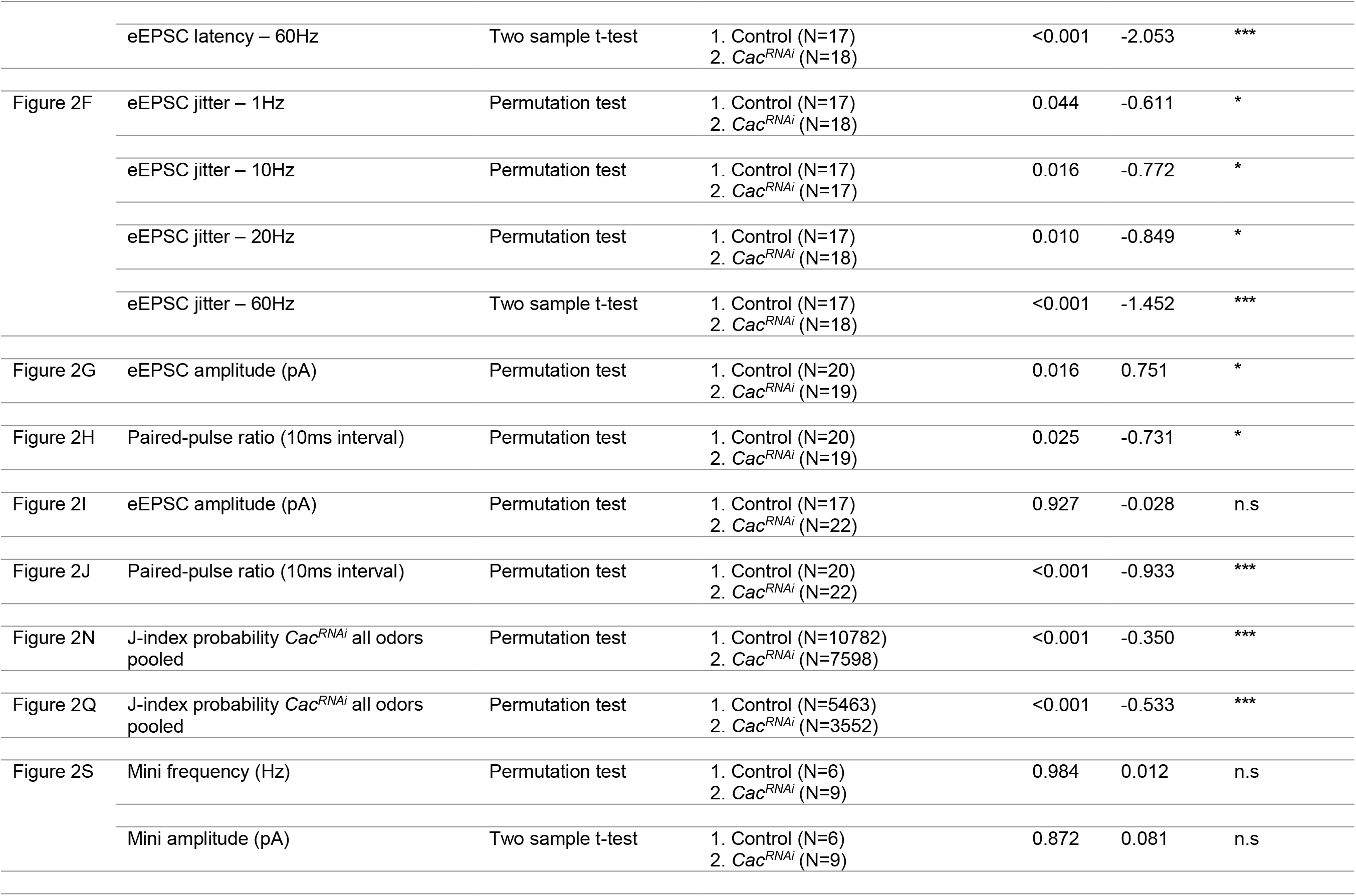
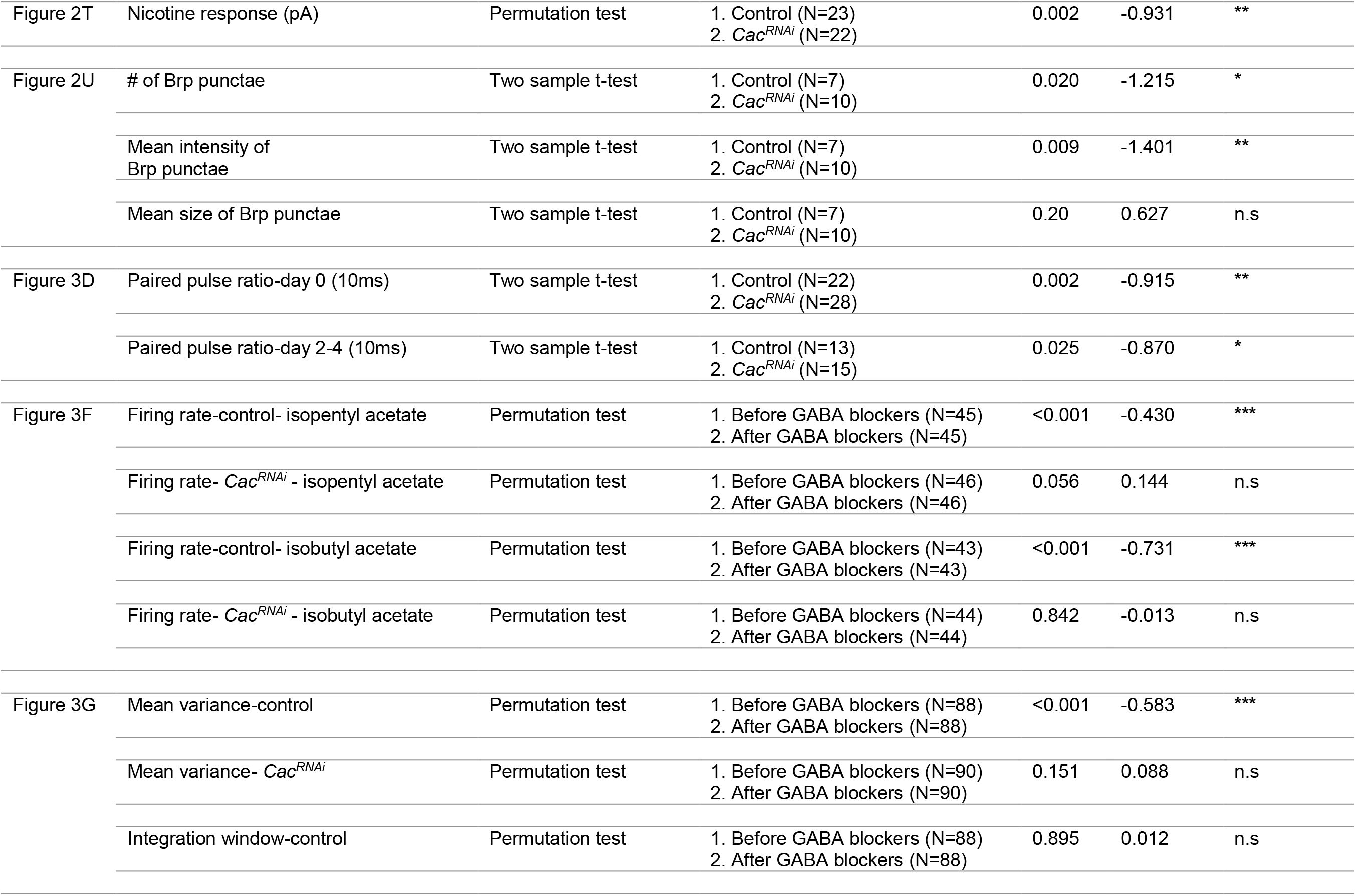
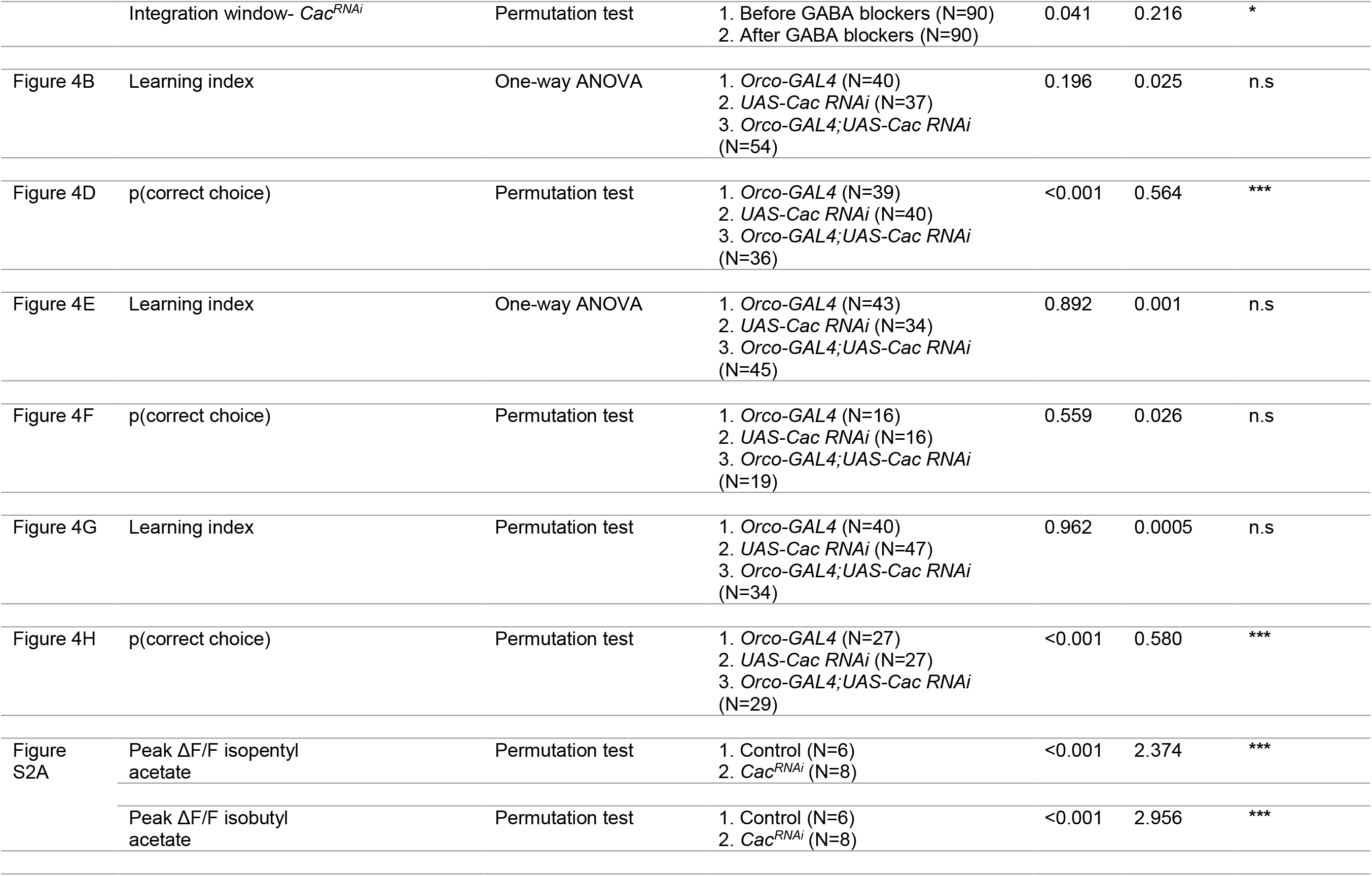
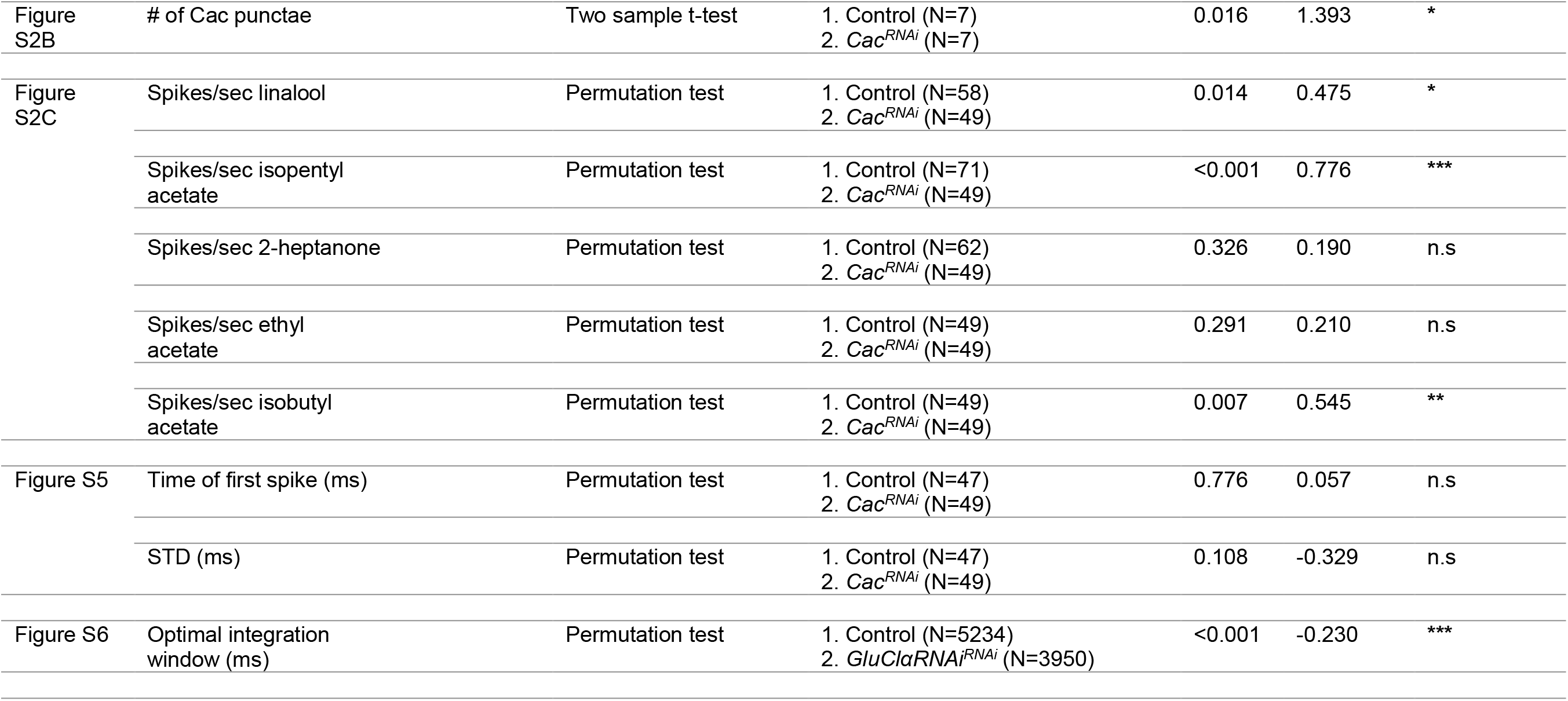
statistical analysis.

